# Reward actively engages both implicit and explicit components in dual force field adaptation

**DOI:** 10.1101/2023.08.09.552587

**Authors:** Marion Forano, David W. Franklin

## Abstract

Motor learning occurs through multiple mechanisms, including unsupervised, supervised (error-based) and reinforcement (reward-based) learning. Although studies have shown that reward leads to an overall better motor adaptation, the specific processes by which reward influences adaptation are still unclear. Here, we examine how the presence of reward affects dual-adaptation to novel dynamics, and distinguish its influence on implicit and explicit learning. Participants adapted to two opposing force fields in an adaptation/de-adaptation/error-clamp paradigm, where five levels of reward (a score and a digital face) were provided as participants reduced their lateral error. Both reward and control (no reward provided) groups simultaneously adapted to both opposing force fields, exhibiting a similar final level of adaptation, which was primarily implicit. Triple-rate models fit to the adaptation process found higher learning rates in the fast and slow processes, and a slightly increased fast retention rate for the reward group. While differences in the slow learning rate were only driven by implicit learning, the large difference in the fast learning rate was mainly explicit. Overall, we confirm previous work showing that reward increases learning rates, extending this to dual-adaptation experiments, and demonstrating that reward influences both implicit and explicit adaptation. Specifically, we show that reward acts primarily explicitly on the fast learning rate and implicitly on the slow learning rates.

**New and Noteworthy:** Here we show that rewarding participants’ performance during dual force field adaptation primarily affects the initial rate of learning and the early timescales of adaptation, with little effect on the final adaptation level. However, reward affects both explicit and implicit components of adaptation. While the learning rate of the slow process is increased implicitly, the fast learning and retention rates are increased through both implicit components and the use of explicit strategies.

## Introduction

Humans continually adapt to both internal and environmental changes to produce appropriate and accurate movements. In this process called motor learning, humans form motor memories consisting of internal models representing the body and the environment. To maintain a high level of motor performance, these internal models are continuously updated, using current environment and body states to adjust the next movement in order to reduce their movement error. Therefore, this error-driven learning requires the use of sensory prediction errors to update the next motor command. While motor learning is primarily error-driven, additional or alternative information can be used. For example, external information, such as positive (reward) or negative (punishment) reinforcers, can be triggered by the current movement. In this case, humans would change their movement, or adapt, in order to maximise the overall task success [1].

This process, called reinforcement learning, has been applied in motor adaptation and is important for understanding underlying processes of motor memory formation [2–8]. This method has been mainly used with a reward, or positive conditional reinforcer, defined as a stimulus administered to an organism following a correct or desired response that increases the probability of occurrence of the response [9]. In general, the presence or absence of a reward is compared to the expectation of such a reward to estimate the reward prediction error which drives changes in behavior. If error feedback is present in the current state, the sensorimotor control system would use sensory prediction errors and reward prediction errors. On the other hand, in the absence of error information, it would only rely on reward prediction errors to estimate the correct or desired response. Recently it has been suggested that rewards and punishments act at a meta-learning level, directly controlling the learning rates to maximize the reward or minimize the punishment [10]. While reinforcement learning has already been extensively studied in many fields, particularly in both psychology and robotics, fewer studies have investigated its effects in motor adaptation.

Most studies investigating reinforcement learning in motor adaptation, have shown that the presence of reward generally leads to stronger motor learning [6, 11–13], although this varies in terms of the effects. This variability in specific effects likely arises, both due to the variability in the type of motor adaptation task (serial reaction time, visuomotor rotation, or force field adaptation) and the type of reward provided (visual, monetary or praise). For example, one study has shown that reward increases both the learning and retention rates for stroke patients in force field adaptation [14]. However, another study that used visual reward during visuomotor rotation, has argued that reward only increases the learning rate, producing faster learning [13]. In contrast, several studies have suggested that reward mainly enhances adaptation through a greater retention rate, for a range of tasks including isometric tracking [15], sequence learning [16] or visuomotor rotation on healthy [17–20], cerebellar degenerative patients [21] or stroke patients [22].

Studies on visuomotor rotation offer strong evidence that visuomotor adaptation is driven primarily by explicit adaptation [23–31]. Additionally, they revealed that reward plays an important role in the use of these explicit strategies [32–34]. In their study, Codol and collaborators [32] demonstrated that the use of explicit strategies in rewarded visuomotor adaptation enhances the retention of adaptation and that the prevention of these aiming strategies reduces motor adaptation. Similarly, Holland and collaborators [33] showed that reward-based gradual-adaptation to a visuomotor rotation is mainly driven by an explicit component, which can also prevent learning when the task becomes more complex, here with the addition of a mental rotation task.

Only a few motor adaptation studies have examined the effect of reward on the learning of novel dynamics, showing its importance in forming and retaining motor memories with a binary reward [35], different monetary reward probability distributions [36], or through performance-scaled reward on stroke patients [14]. While there is strong evidence that visuomotor adaptation is driven primarily by explicit adaptation, recent studies have suggested that force field adaptation is primarily implicit in nature [37, 38]. Due to these different natures, one could imagine that reward has different effects on each type of learning task. Moreover, both force studies examining reward on force field adaptation tested the adaptation to only one set of dynamics at once. That is, they tested single adaptation to a force field. However, in daily life, humans continuously switch from one tasks to another, often requiring different internal models to deal with different external dynamics. Such adaptation can be examined using a dual-adaptation paradigm in which participants adapt simultaneously to two different dynamics, each signalled by a contextual cue.

When appropriate contextual cues are provided, humans are able to select, recall and adapt the respective model memory for each task [38–46]. Dual-adaptation has been shown to occur when the contextual cue is different in the physical or visual hand state [42, 43], workspace visual location [43, 47], bimanual or unimanual arm movements [40, 48], some body postures [49] or for lead-in or follow-through movements [41, 50–52]. In contrast, other cues such as color provides only a weak or ineffective contextual cue for dual-adaptation [43, 53], despite some studies showing a small effect [54–56]. These differences in the efficiency of the color cues has recently been explained by the use of explicit strategies when the task is sufficiently simple [38]. Specifically, they showed that dual-adaptation occurs implicitly when a direct contextual cue is used (a contextual cue related to the physical or visual hand state) [38]. However, participants apply explicit strategies in simple tasks when the contextual cues relate to more abstract representations (indirect contextual cues); for example a difference in background, target or cursor color. The differences in the effects of indirect and direct cues on dual-adaptation, and the presence of explicit and implicit components during force field adaptation, raise critical questions about the effect of reward on motor adaptation, particularly within a force field dual-adaptation task. As found in visuomotor rotation studies, we predict that reward strongly influences explicit adaptation. Here, we examine whether reward affects the adaptation to two opposing force fields, and to what degree any differences occurs within the implicit or explicit components of motor adaptation. To address these questions, we designed a force field experiment including visual rewards based on participants’ performance to a reaching task. Five different types of rewards were scaled to the participants’ lateral error such that the visual reward got stronger as participants reduced their error. An adaptation/de-adaptation/error-clamp paradigm was used in order to facilitate the interpretation in comparing the results to previous studies [45, 57]. Here we propose that rewarding participants’ performance leads to greater dual-adaptation, through better retention and possibly faster learning. Additionally, we investigate the possibility of this increase in adaptation arising through explicit components.

## Methods

### Participants

A total of 28 (16 male, 12 female, mean age 31.1 *±*1.1 years) force field naive participants, with no known neurological disorders, participated in the experiment. All participants were right-handed based on the Edinburgh handedness questionnaire [58] and provided written informed consent before participation. The study was approved by the Ethics Committee of the Medical Faculty of the Technical University of Munich.

### Apparatus

Seated in a custom adjustable chair and strapped with a four-point seat belt to reduce body movement, participants grasped the endpoint handle of a vBOT robotic manipulandum [59], with their forearm supported against gravity by an air sled. The vBOT system is a custom built robotic interface that can apply state-dependent forces on the handle while recording the position and velocity in the planar workspace, located approximately 15 cm below the shoulder. A six-axis force transducer (ATI Nano 25; ATI Industrial Automation) measured the end-point forces applied by the participant on the handle. Joint position sensors (58SA; Industrial encoders design) on the motor axes were used to calculate the position of the vBOT handle. Position and force data were sampled at 1kHz. Visual feedback to the participants was provided horizontally from a computer monitor (Apple 30” Cinema HD Display, Apple Computer, Cupertino, CA, USA; response time: 16 ms; resolution: 2560 *×* 1600) fixed above the plane of movement and reflected via a mirror system that prevented visual feedback of the participants’ arm. Constant visual feedback, provided in the same plane as the movement, consisted of circles indicating the start, target and cursor positions on a black background (Fig 1A). Necessary auditory feedback was provided by speakers. Specific visual and auditory feedback is defined in the experimental paradigm.

**Figure 1.**
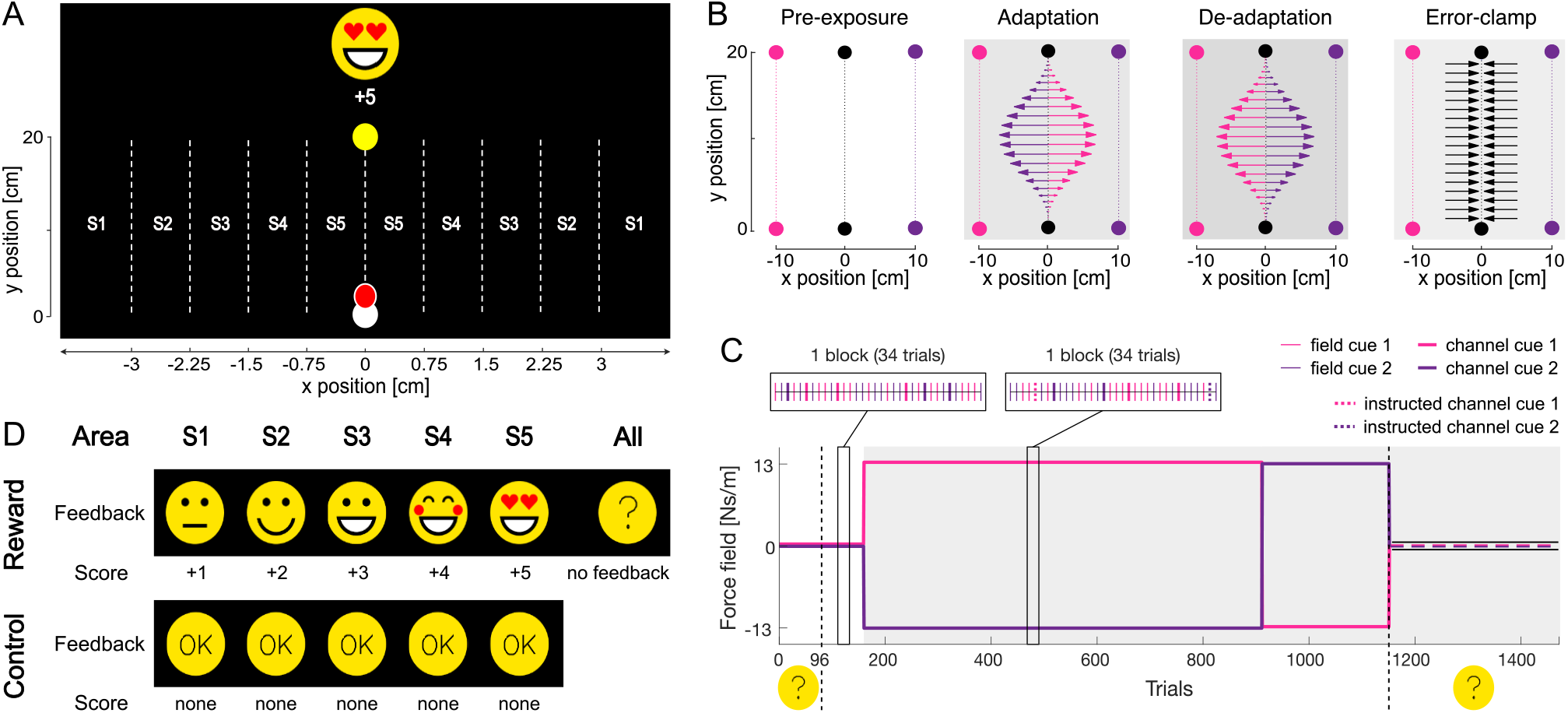
Experimental Setup and Paradigm. A. Different areas of the workspace were divided to provide different reward regions depending on the maximum lateral error (x-axis), which are indicated between S1 and S5. B. Workspace layout of the experiment displaying the four different phases with their respective force fields and cues. In the adaptation phase, two force fields were applied (CW and CCW), where each force field was always associated with one of the contextual cues (e.g. CW force field for the left visual workspace and CCW force field for the right visual workspace). In the de-adaptation phase, the association of the force fields to the visual cues was reversed (e.g. CCW force field for the left visual workspace and CW force field for the right visual workspace. Participants always physically performed forward reaching movements in the center of the workspace (black) while visual feedback (targets and cursor) was presented at one of the two visual workspaces which acted as a contextual cue:-10 cm offset (left workspace, pink) set as cue 1 and +10 cm offset (right workspace, purple) set as cue 2. Pink and purple colors are only used here for illustration. C. Temporal structure of the experiment. An adaptation (middle grey)/de-adaptation (dark grey)/error-clamp (light grey) paradigm was performed over a maximum of 1632 trials. In the error-clamp phase (light grey), the kinematic error is held at zero to assess spontaneous recovery. The structure of pseudo-randomized blocks of trials are displayed for the pre-exposure (field and normal channel trials) and adaptation phase (field, normal channel and instructed channel trials). Instructed trials were only used during the adaptation and the de-adaptation phase. Additional “hidden reward” feedback phases are displayed, happening for the first half of the pre-exposure phase and all the error-clamp phase only. D. Visual feedback and score grading. Reward participants experience different visual rewards scaled from the area S1 to the area S5. A reward was composed of both a increase in points and the presentation of a digital face. Control participants were given a neutral yellow circle with a centered “OK” when they reached the speed requirement and were not provided with any points.

### Experimental setup

Participants made right-handed forward reaching movements from a start position (grey circle of 1.5 cm diameter) located approximately 20 cm directly in front of the participant, to a target (yellow circle of 1.5 cm diameter) located 20 cm away to the front (Fig 1A). A trial was initiated by the cursor entering the start position. The go-cue was defined as the target appearing after the cursor resided in the start position for a random exponentially distributed time between 1 and 2 s. The movement was considered complete when the participants maintained the cursor within the target for 600ms. After each trial, visual feedback regarding the success of the previous trial was provided while the participant’s hand was passively moved back to the start position by the robotic manipulandum. Successful trials were defined as trials where participants hit the target without overshooting and where their movement’s peak speed was between 37 and 53 cm/s. On these trials, the participants received specific feedback, defined below, depending on the group they belonged to. On unsuccessful trials, messages of “too fast”, “too slow” or “overshoot” were provided when the peak speed exceeded 53 cm/s, did not reach 37 cm/s or when the cursor overshot the target by more than 1 cm, respectively. An additional low tone was played on these trials. If participants failed to leave the start position within 1000 ms after the target appearance, the current trial was aborted and restarted. Short breaks were enforced every 200 trials except for the first break that was set after 140 trials, in order to repeat specific instructions before exposure to force fields. Movements were self paced, and participants were also able to take a short break before starting the next trial if desired. Participants were instructed to produce one powerful reaching movement to the target as soon as the target appeared (go-cue). During each movement, the vBOT was either passive (null field), produced a clockwise (CW) or counterclockwise (CCW) velocity-dependent curl force field, or a mechanical channel (Fig 1B-C). For the velocity-dependent curl field [53], the force at the handle was given by:

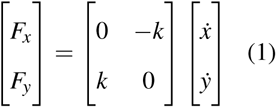

where k was set to either *±*13 *N · m^−^*^1^ *· s*, with the sign of k determining the force field direction (CW or CCW). On a channel trial, the reaching movement was confined to a simulated mechanical channel with a stiffness of 6000 *N · m^−^*^1^ *· s* and damping of 2 *N · m^−^*^1^ *· s* acting lateral to the line from the start to the target [60, 61]. These “channel” trials were used to measure the lateral force, produced by the participants on the wall of this mechanical channel, that reflects their compensation to the exposed force field. “Instructed channel” trials were similar to channel trials but were preceded by the audio message “Robot off” [38], informing participants that the environmental disturbance was removed for this specific trial only. These instructed trials allowed us to assess the amount of implicit adaptation employed by participants [37, 38]. Here, by defining the total amount of adaptation as the summation between the explicit and implicit components, we can calculate the amount of explicit strategies by taking the difference between the total amount of adaptation and the amount of implicit adaptation [62, 63].

### Experimental Paradigm

All participants were required to adapt to an adaptation/de-adaptation/error-clamp, or A/B/error-clamp, paradigm [57] within dual-adaptation [45]. In this paradigm, after a pre-exposure phase containing 5 blocks of trials (each block contained 34 trials), participants were simultaneously presented with two opposing force fields in an adaptation phase (exposure to fields A), containing 28 blocks, and a consecutive de-adaptation phase (exposure to reversed fields B) containing between 2 and 10 blocks of trials depending on the participants’ performance, followed by a final error-clamp phase, containing 5 blocks of trials (Fig 1B-C). To allow dual-adaptation, each of the two opposing force fields was linked with an appropriate contextual cue: a shift in the workspace visual location [38, 43] either to the left (−10 cm from the sagittal axis, Fig 1B pink workspace) or to the right (+10 cm from the sagittal axis, Fig 1B purple workspace) of the screen. The physical hand location (proprioceptive location) remained centered for both cues, without any shift from the sagittal axis (Fig 1B black workspace). In the pre-exposure phase, movements with both contextual cues occurred in the null field. Within the adaptation phase one contextual cue (e.g. +10 cm visual shift) was associated with one force field (e.g. CW), whereas the other contextual cue (e.g.-10 cm visual shift) was associated with the other force field (e.g. CCW) (Fig 1B-C). In the de-adaptation phase, the cue-field association was reversed (e.g. CCW for the +10 cm visual shift and CW for the-10 cm visual shift) to drive de-adaptation (Fig 1B-C). The length of the de-adaptation phase was dependent on each participant’s performance and could vary between 2 and 10 blocks of trials (68–340 trials), as detailed in [45]. Specifically, the mean of force compensation over the last three blocks of trials was calculated online for each contextual cue. Once the difference in mean of the two contextual cues switched sign (became negative), which would represent the force compensation of one cue crossing the other, the phase was terminated and switched to the following error-clamp phase, after the current block of trials. Finally, the error-clamp phase exclusively applied channel trials regardless of the current cue. One block of trials was pseudo-randomly composed of 34 trials: 17 trials for each cue including 14 normal trials, and 3 channel trials. During the adaptation and de-adaptation phase, one out of 3 channel trials was replaced by an instructed channel trials (Fig1C), such that one block of trials included 14 normal trials, 2 channel trials and 1 instructed channel trial per cue. While normal trials were presented in a null, CW or CCW force field depending on the temporal phase, channel and instructed channel trials remained error-clamped.

Participants were separated into two groups and either presented with the reward condition or the control condition. In the reward group (8 male, 6 female, mean age 32.7 *±* 1.6 years), participants were presented with a reward for each successful trial. Five different rewards, consisting of five different scores and digital faces, were designed depending on the current trial’s kinematic error. These consisted of a “neutral” (+1 point in score), “satisfied” (+2), “very satisfied” (+3), “happy” (+4), and “very happy” (+5) digital face (Fig 1D). The face was continuously presented as a 6 cm-diameter yellow circle on top of the screen (Fig 1A), with its face properties (eyes, mouth, cheeks) changing and transferring different “emotions” depending on the kinematic error. Specifically, the maximum perpendicular error on each trial was used to determine the level of reward given to the participants in the reward group. This measure reflected participants’ performance on each trial. We present feedback on the kinematic measure of each trial rather than only for force compensation on the channel trials, as these are most frequently encountered by participants and allow the presentation of reward feedback as often as possible. It is important to note that participants can reduce the kinematic error to increase reward through predictive compensation to the force field, through increased co-contraction, limb stiffness and feedback gains, or through combinations of all of these [64, 65]. Five ranges of *±*0.75 cm error were defined to provide the different rewards (Fig 1B). The first closest range to the perfect straight reaching movement (Fig 1A,D S5, lower error possible, <0.75 from the target) to the furthest range (S1, higher error possible, >3 cm from the target) were respectively linked to the reward scale from “very happy” (S5, +5 points in score, highest reward possible) to “neutral” (S1, +1 point in score, lowest reward possible). Rewards were given during the whole experiment except during the first half of the pre-exposure phase and the full error-clamp phase. In these “hidden reward” phases, the yellow digital face only displayed a centered question mark (Fig 1C) and the score was frozen. However, participants were still asked to perform accurate movements, regardless of the phase. Specifically, in the “hidden reward” error-clamp phase, while the displayed score was frozen, the hidden total score took into account the points gathered and was finally displayed at the end of the experiment. For unsuccessful trials, no reward (+0 points in score) was given to the participants, causing the digital face to disappear until the next successful trial.

In the control group (8 male, 6 female, mean age 31.6 *±* 2.0 years), no reward was given throughout the experiment. However, to differentiate successful from unsuccessful trials, a similar 6 cm-diameter yellow circle with a centered “OK” was continuously presented on top of the screen (Fig 1D) when participants succeeded, and disappeared when participants failed to have a successful trial. No score was given to participants in this control condition.

Each group (condition) was counterbalanced such that half of the participants experienced the adaptation phase with contextual cues matched to one set of force field directions, whereas the other half of the participants experienced contextual cues matched to the opposite force field directions.

### Analysis

Data were analyzed offline using MATLAB (R2022a, The MathWorks, Natick, MA). All measurements were low-pass filtered at 40 Hz using a 10^th^ order zero-phase-lag Butterworth filter (filtfilt). Over the course of the experiment, the lateral force measured can vary due to the natural drift of the mass of the arm over the air-sled. In order to avoid interference in our measurements from this drift, we subtracted the mean offset in lateral force for each participant measured between 250 and 150 ms prior to the movement start, from the total force trace. The start of the reaching movement was defined as the cursor leaving the start position (cursor center crossing the start position’s 1.5 cm-radius) and the end was defined as the cursor entering the target (cursor center crossing the target’s 1.5 cm-radius). In order to quantify adaptation, the following measures were calculated.

#### Success

On field trials (null or curl field trials), we defined participants’ success as both their success type on each trials and their level of overall percentage of success. The success type was defined as the rewarding score given for the current trial and the level was defined as the mean across blocks of trials and calculated for plotting purposes. The percentage of success was defined as the percentage of each given success’ type (0, 1, 2, 3, 4 and 5 separately) over all the experiment. As participants in the control group were not provided any rewards, we estimated offline the success level equivalent to their performance and compared them to the given success level of the reward group.

#### Trajectory bias and variability

We calculated both bias and variability in trajectories to explore differences in movements between the control and the reward groups, and their consistency over time. We first divided each reaching movement into 100 evenly spaced data points between the start and the end targets. For each of these 100 data points, we calculated the mean and standard deviation across the last five blocks of trials for the pre-exposure and adaptation (late adaptation). Additionally, to explore the evolution of any biases over time, the mean bias across the five blocks of trials of pre-exposure, the first block of adaptation (early adaptation), the last five blocks of adaptation (late adaptation), and the last two blocks of de-adaptation (late de-adaptation) were calculated. The mean and standard-error of the mean across participants were then calculated for each of these data points, in order to display the trajectories. The variability was finally calculated for each participant by averaging the standard deviation of the trajectories (100 data points per trajectory) across both pre-exposure and late adaptation stages.

#### Kinematic error

For each field trial (null or curl field depending on the temporal phase), the maximum perpendicular error (MPE) was calculated and used as a measure of kinematic error. MPE is defined as the signed maximum perpendicular distance between the hand trajectory and the straight line joining the start position and the current displayed target.

#### Force compensation

For both channel and instructed channel trials, the force compensation was calculated to quantify the amount of predictive force adaptation to the force field. Force compensation is defined on each trial by regressing the end-point force applied by the participant on the handle (lateral measured force) against the force needed to compensate perfectly for the force field [57]. Specifically, the slope of the linear regression through zero is used as a measure of force compensation. The perfect compensatory force was determined as the forward velocity of the current trial multiplied by the force field constant k. As each group was counterbalanced across participants, the values were flipped such that the force compensation associated with each cue had the same sign for all participants. Since the measure of force compensation is relative to the field, we inverted one cue for plotting in order to visually differentiate the two opposing forces. All models used to fit the data assume equal adaptation to both force fields. In order to allow this equal adaptation, we subtracted block-wise the mean across contextual cues from the force compensation values for each contextual cue (Fig 4B).

#### Relative lateral force

To examine the shape and timing of the force applied by the participant to compensate for the disturbance, the relative lateral forces were calculated. Individual force trials were aligned to peak velocity, and clipped between-300 and +300 ms from this peak. For averaging and visualization, we normalized forces in the x-axis by the peak of the perfect compensation force profile. This perfect force profile was calculated as the y-axis velocity multiplied by the force field constant k.

#### De-adaptation phase

Throughout the experiments, data are primarily presented for each block. However, participants in the de-adaptation phase performed a different number of trials (blocks). This phase was divided into equal-sized sections, where the mean data was determined for each section rather than for each block in order to allow averaging across participants. 35 (control) and 14 (reward) averaging sections were used for kinematic error, whereas 10 (control) and 4 (reward) averaging sections were used for force compensation.

### Model fitting

#### Adaptation Rate

In order to examine differences in the trial time constant of adaptation and final asymptote of the initial adaptation phase we fitted an exponential function to the force compensation for each trial throughout this phase of the experiment. Here we used the force compensation of both cues, such that the force compensation for each cue increased over the adaptation towards positive adaptation (both cues towards 100%), and could reflect the total adaptation. We found the parameter values that best fit the experimental data, using a least-squares cost function (fminsearchbnd), where the asymptote parameters were constrained between 0 and 100, and the trial time constant was constrained between 0 and 10. The adaptation rate and asymptote parameters were estimated using a leave-two out cross-validation sampling method to obtain appropriate significance levels. A total of 91 sets of parameters were obtained from all sample combinations of 12 out of 14 participants data. The optimization was performed ten times for each sample, with a random initial parameter setting within the parameter constraints, and the one with the smallest sum of squares error was used. For both groups, the force compensation of normal and instructed channel trials were used as two separate input variables of the fitting, resulting in independent parameter sets.

#### Learning and retention rates of the multi-rate model

Participants’ force compensation over the entire experiment for was fitted with a weighted triple-rate model [45, 66]. This model assumes that adaptation occurs through the summation (and competition) of two separate states weighted by contextual cues (**c**), at a fast (f), slow (s) and ultra-slow (us) timescale of learning and retention. Therefore, one set of parameters included a retention rate *A* and a learning rate *B* for a fast, slow and ultra-slow process, as well as one overall weighted switch *C*.

The model we used in order to fit experimental data is defined as

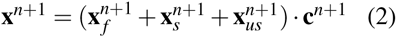

where

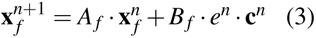

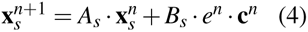

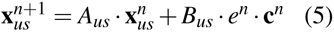

and

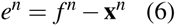

with

**x***^n^*^+1^ – output on subsequent trial

**x***^n^* – output on current trial

*e^n^* – error on current trial

*A* – retention rate

*B* – learning rate

*f ^n^* – environmental force

c = [C 1 −C] where C is set as the weighted switch estimated parameter

First, the mean of force compensation values over the two cues was subtracted to remove any potential bias between the cues. Then, the force compensation was fitted with the model using a least-squares cost function (fminsearchbnd). We use a leave-two out cross-validation sampling method in order to obtain appropriate significance levels. For this, we estimate each set of parameters by fitting the model to the force compensation of a sample of 12 participants. A total of 91 sets of parameters were obtained from all sample combinations of 12 out of 14 participants data. To obtain each of these 91 sample combinations, the optimization was performed five times using a random initial parameter setting within the parameter constraints (see below). Out of these five fits, the parameter set that resulted in the smallest error, was kept as the single parameter set for this specific sample combination. The random initial parameters were constrained as following:

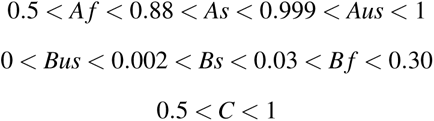

In order to compare both groups and adaptation components, we fitted the force compensation of normal and instructed channel trials as two separate input variables, for both the control and reward group. Therefore, this model parameter optimization process resulted in four batches of 91 sets of parameters: one batch of 91 parameters sets for the Control normal group, one batch for the Reward normal group, one batch for the Control instructed group and one batch for the the Reward instructed group. Only essential comparisons were analyzed such that the parameters of the Reward instructed trials (Ri) were not compared with the Control normal trials (C) as well as the Control instructed trials (Ci) with the Reward normal trials (R).

### Statistics

All statistical tests below were performed using JASP Team (2022), JASP (Version 0.16.3)[computer software], except for the Kruskal-Wallis tests on the parameter estimates and the t-test on trials number in the de-adaptation phase, that were performed on Matlab (R2022a, The MathWorks, Natick, MA). Further differences between levels were examined using Bonferroni post-hoc comparisons after each Analysis of Variance (ANOVA).

#### Success

T-tests, were performed for all comparisons between the reward and control groups, except for the success type 0 where a Mann-Whitney t-test and for the success type 4 and 5 where a Welch’s t-test were performed instead, due to the inequality of variances.

#### Variability

A repeated measures ANOVA with a main effect of stage (2 levels: pre-exposure and late adaptation) and the between-subject factor group (2 levels: control and reward) was performed in order to compare the variability in the pre-exposure and late adaptation stages between groups.

#### Kinematic error and force compensation

For the kinematic error, a repeated measures ANOVA with a main effect of stage (4 levels: pre-exposure (all blocks of trials), early adaptation (first block of trials), late adaptation (last five blocks of trials) and late de-adaptation (last block of trials)) and the between-subject factor group (2 levels: control and reward) was used. For comparing the total adaptation between groups and stages, we applied a similar repeated measures ANOVA on the force compensation on normal channel trials, with a main effect of stage (4 levels: pre-exposure (all blocks of trials), late adaptation (last five blocks of trials), late de-adaptation (last block of trials) and error-clamp (all blocks of trials)) and the between-subject factor group (2 levels: control and reward). A Greenhouse-Geisser sphericity correction was used on the within-subject effect for both the kinematic error and the force compensation. All post-hoc comparisons were performed using a Bonferroni correction. Additionally for the error-clamp phase, the difference in force compensation between both cues was calculated and compared to zero using a one-sample t-test for the control group and a one-sample Wilcoxon test for the reward group. To compare the implicit and total adaptation, we performed two-samples Wilcoxon tests between the two channel types in each group for the end stage of adaptation (5 last blocks) and de-adaptation (3 last blocks), except for the end stage of adaptation in the Reward group where a student t-test was performed.

#### Trials number difference

In order to compare the length of de-adaptation between the control and the reward group, we performed a two-sample t-test using Matlab on the number of trials for each participants across groups.

#### Retention and learning rates

To investigate the differences in retention and learning rates between the control and reward groups, as well as the normal and instructed trial types, we performed Kruskal-Wallis tests on the parameter estimates: 91 samples in each of the four data sets (C, Ci, R, Ri) for all seven parameters (Af, As, Aus, Bf, Bs, Bus, Switch parameter C). Differences that were found at a level below under the 7th-digit were considered invalid due to the accuracy of floating point values in the software (Matlab R2022a). Therefore, we did not consider any of their statistical effect.

## Results

Here, we investigated the influence of reward on dual-adaptation to novel force fields. One group of participants was given different levels of reward depending on their lateral kinematic error (reward group), while another group received no additional feedback about their lateral error (control group). Both groups were provided with online visual feedback of their movements. Both groups adapted to two opposing force fields (separated with visual offsets cues) using an adaptation/de-adaptation/error-clamp paradigm, allowing us to examine the learning and retention parameters of adaptation and compare differences in the formation of motor memories with and without reward.

### Success

What differentiated the reward and control groups is the inclusion of reward, comprised of both visual feedback and scores ordered in five different sections (from type 1, lowest reward, to type 5, highest reward) for the reward group, which did not exist for the control group. We examined the effect that this feedback had on the specific levels of success obtained by the participants across the experiment (Fig 2A). For the reward group, this success type was provided directly to the participants after each trial, whereas for the control group this success type was calculated but never provided to the participants. Although similar in the pre-exposure phase across groups (t_26_=1.154; p=0.259), the mean success level increased faster (2A) with reward during adaptation, with evidence of a higher final level than the control group (t_26_=-2.510; p=0.019). Similar results were also seen in the de-adaptation phase, with faster increases in success and higher final levels for the reward group (t_26_=-3.128; p=0.004). Note that a trial was considered successful if it initially satisfied movement speed requirements (otherwise the success was set as 0) and was additionally assigned a score (between 1 and 5) depending on its lateral kinematic error (maximal perpendicular error). Importantly we found no difference in the percentage of unsuccessful trials (Fig 2B) between the control (31.6%) and reward (29.2%) groups (U=97.000; p=0.982) across the whole experiment. This implies that the presence of the reward feedback did not influence the overall success or failure rate (for example through reducing the speed of movements to better counter the disturbance) but appears to primarily have an effect on the relative level of success, through reductions in the lateral error.

**Figure 2.**
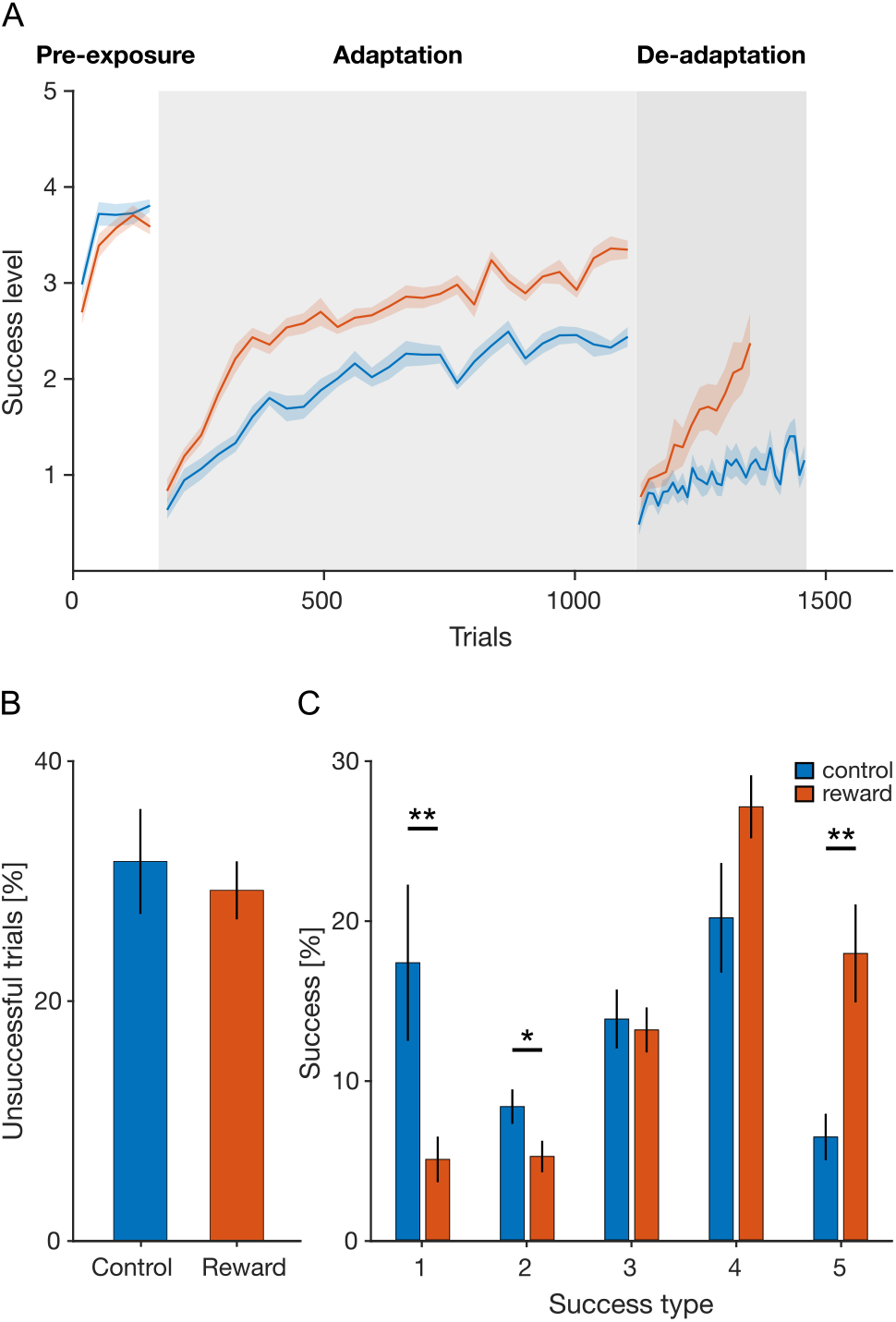
Success level. A. Mean success level for the control group (blue) and reward group (dark orange) during the pre-exposure, adaptation and de-adaptation phases (normal trials only). The mean (solid line) and standard-error (shaded area) of the success level across participants was calculated for each block. B. Mean and standard-error of the percentage of unsuccessful trials (success type 0) during the pre-exposure, adaptation and de-adaptation phases. C. Mean and standard-error of the percentage of successful trials for the 5 different success levels across the first three phases. Channel trials are excluded in this analysis.

Reward feedback had the largest influence on the distribution of success type (Fig 2C). Although participants in both groups obtained similar numbers of trials with success type 3 (t_26_=0.293; p=0.772) and type 4 (t_20.734_=-1.757; p=0.094), the reward participants attained a higher number of trials with success type 5 (t_18.569_=-3.385; p=0.003, with reducing their success type 1 (U=159; p=0.005) and type 2 (t_26_=2.138; p=0.042). The reward group resulted in participants obtaining higher scores through making movements with smaller lateral errors, and this from an early stage in the adaptation phase.

### Trajectory bias and variability

In addition to the effects on success levels, the presence of reward could have an effect on the kinematic trajectory, in particular in the bias and variability of the paths. The bias, or mean trajectory, used to reach the target location (Fig 3A, solid lines) and its variability across participants was examined across four stages of the experiments. As expected, no differences were seen in the pre-exposure stage. It can be seen that the reward group exhibited smaller deviations from the straight line between the start and end targets, particularly in the late adaptation and late de-adaptation stages. In addition, we examined the effect of reward on the variability of trajectories around each participant’s mean in the late adaptation stage (Fig 3B). A repeated measure ANOVA with a main effect of stage (2 levels: pre-exposure and late adaptation) and the between-subject factor group (2 levels: control and reward) showed a main within-subjects effect of stage (F_1,26_=62.065; p<0.001), of the interaction group*stage (F_1,26_=9.512; p=0.005) and for the between-subjects factor group (F_1,26_=4.685; p=0.040). As expected, no difference in variability is seen between groups in the pre-exposure stage (post-hoc comparison: p=1.000), and participants increase their variability from the pre-exposure to the late exposure stage in both the control (post-hoc comparison: p<0.001) and reward groups (post-hoc comparison: p=0.002). Additionally, we found a difference in the variability of the trajectories between the control and the reward groups at the end of adaptation (post-hoc comparison: p=0.003). This shows that the reduction in trajectory deviation in the reward group occurred with a present but significantly smaller increase in the trial-by-trial variability in these trajectories than the control group.

**Figure 3.**
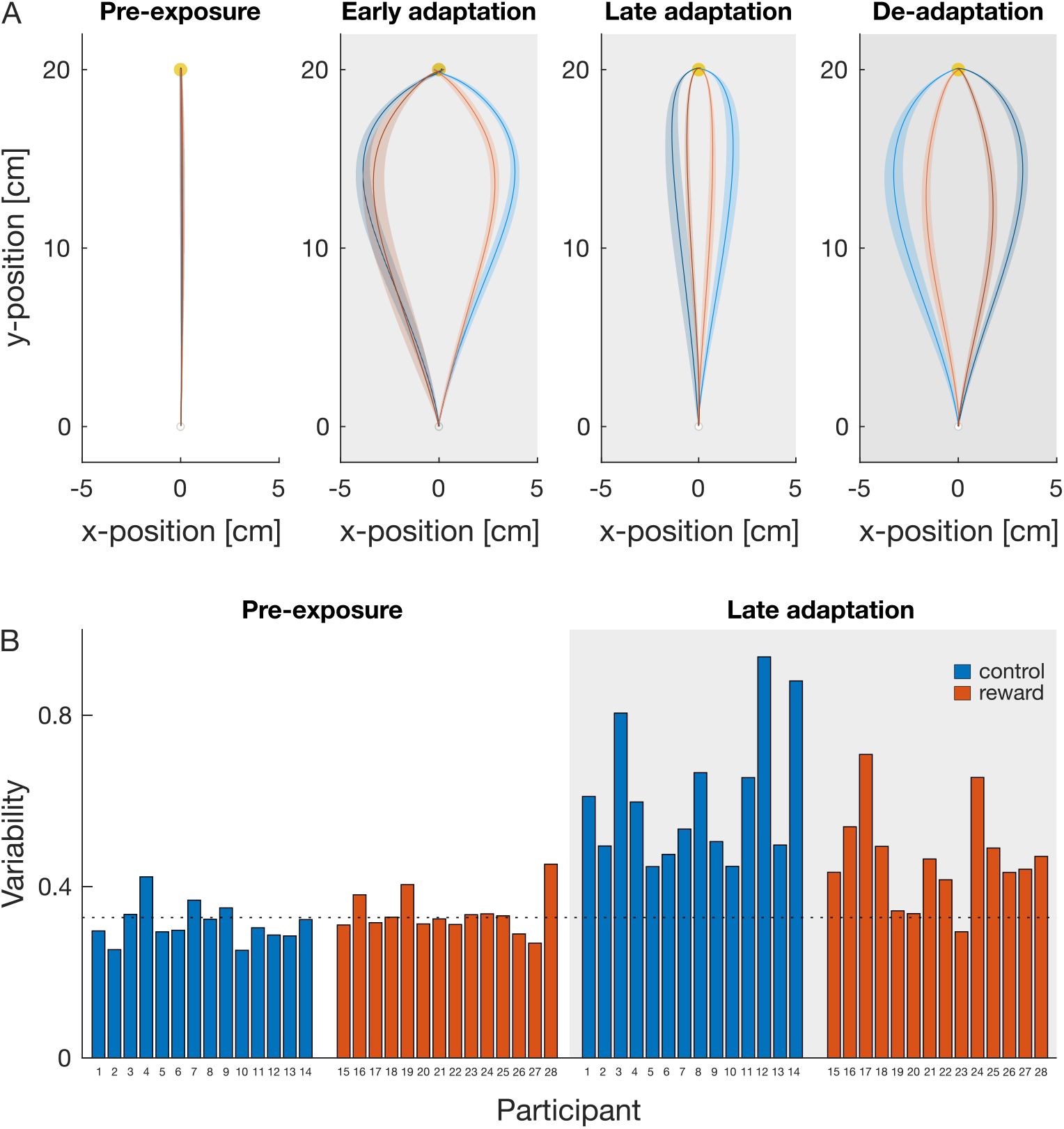
Trajectory bias and variability. A. Mean trajectory (solid line) and standard-error across participants (shaded area) of each cue separately (light and dark solid lines), for the pre-exposure (all blocks), early adaptation (first block), late adaptation (last five blocks) and late de-adaptation (last two blocks) for the control (blue) and reward (orange) groups. B. Mean trajectory variability across pre-exposure (all blocks) and late adaptation (last five blocks) for each participant. The dotted black line represents the mean variability of all participants across both conditions in pre-exposure, and is displayed as a reference. Channel trials are excluded in this analysis.

### Adaptation

Both groups experienced large errors induced by initial force field exposure which were reduced during the adaptation process (Fig 4). Similarly, in the de-adaptation phase, when the association between the force field and the cue was reversed, participants error considerably increased before it reduced back towards the level experienced during the initial exposure (first trials of the adaptation phase). Overall, both groups of participants were able to independently adapt and de-adapt to the opposing force fields, displaying similar end-levels of adaptation but with differences in the rates and extent of de-adaptation (Fig 4C, F) suggesting differences in the underlying processes of adaptation.

**Figure 4.**
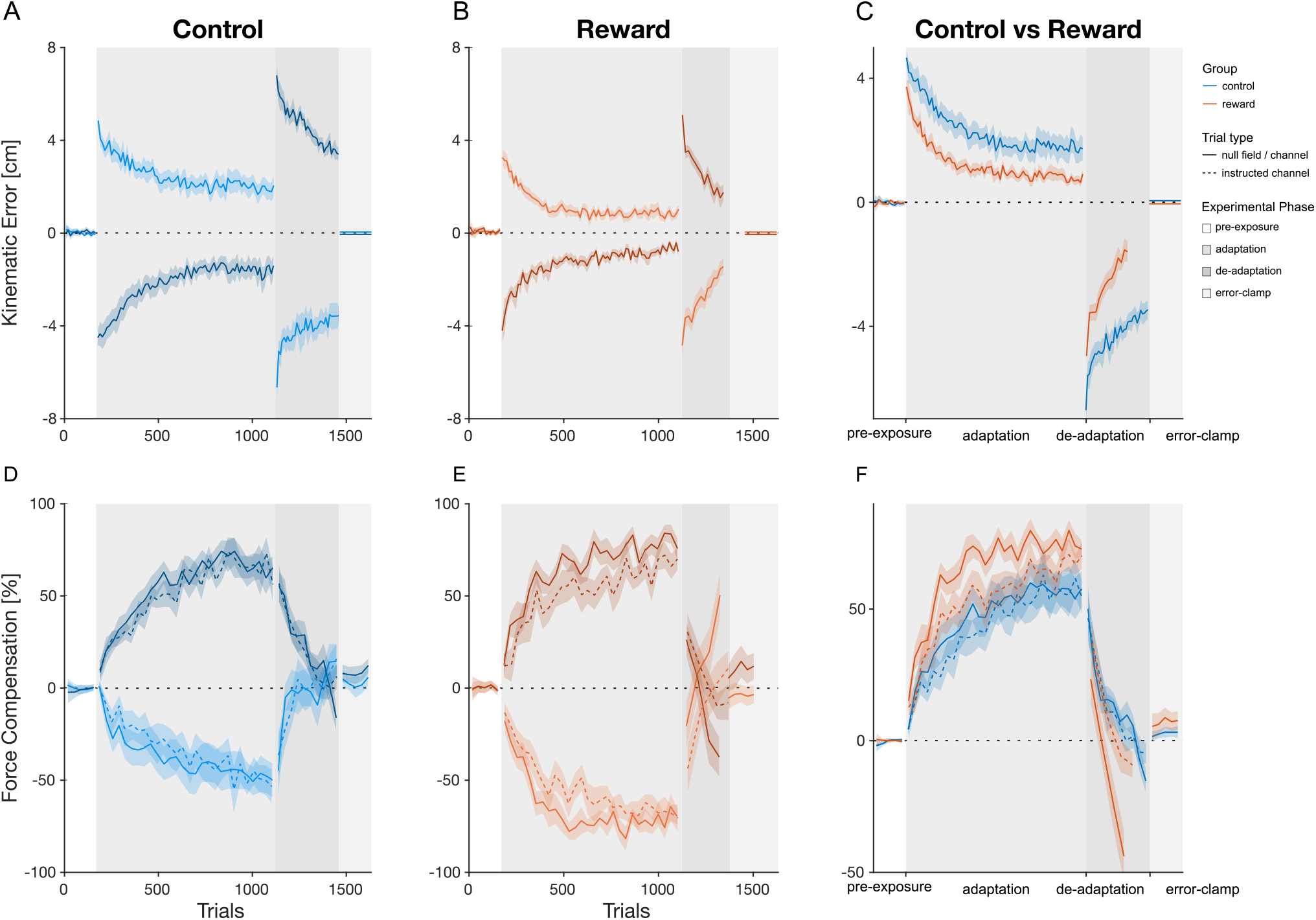
Temporal pattern of adaptation. A-C, Mean kinematic error for both the control (blue) and reward (orange) groups. Mean kinematic error (solid line) and standard error of the mean (shaded region) are shown across the pre-exposure (white), adaptation (grey), de-adaptation (dark grey) and error-clamp (light grey) phases. The data of the contextual cue 1 (left visual workspace shift) and 2 (right visual workspace shift) are presented in light and dark blue (A, control) and orange (B, reward) lines. C. Comparison of the mean across cues between the control (blue) and reward (orange) group. For illustrative purposes the sign of the error was flipped for one of the cues. D-F, Mean force compensation for normal (solid line) and instructed (dashed line) channel trials over the pre-exposure (white), adaptation (grey), de-adaptation (dark grey) and error-clamp (light grey) phases. For illustrative purposes, the data of one of the cues was flipped. The force compensation for the two contextual cues (D,E) is symmetrical due to the subtraction of the mean force compensation across the two cues. F. Comparison of the mean across cues between the control (blue) and reward (orange) group.

We investigated differences in kinematic error using a repeated measures ANOVA with a main effect of stage (4 levels: pre-exposure, early adaptation, late adaptation, late de-adaptation) and the between-subjects factor group (2 levels: control and reward). We found main within-subjects effects of stage (F_1.592,41.391_=181.086; p<0.001) and of the interaction group*stage (F_1.592,41.391_=15.983; p<0.001), but none for the between-subjects factor group (F_1,26_=0.274; p=0.605). The repeated measures ANOVA performed on the force compensation had a main effect of stage (F_1.545,40.171_=128.755; p<0.001) and group*stage (F_1.545,40.171_=9.369; p=0.001), but similarly no evidence of an effect of group (F_1,26_=0.484; p=0.493). Further differences between levels were examined using Bonferroni post-hoc comparisons.

Participants of both groups had the same performance level at the beginning of the experiment (pre-exposure phase, Fig 4), with no differences in kinematic error (p=1.000) or force compensation (p=1.000). Additionally, the relative force profile (Fig 5A) for the two contextual cues in each group were indistinguishable, with force profiles close to the zero level of adaptation as expected.

**Figure 5.**
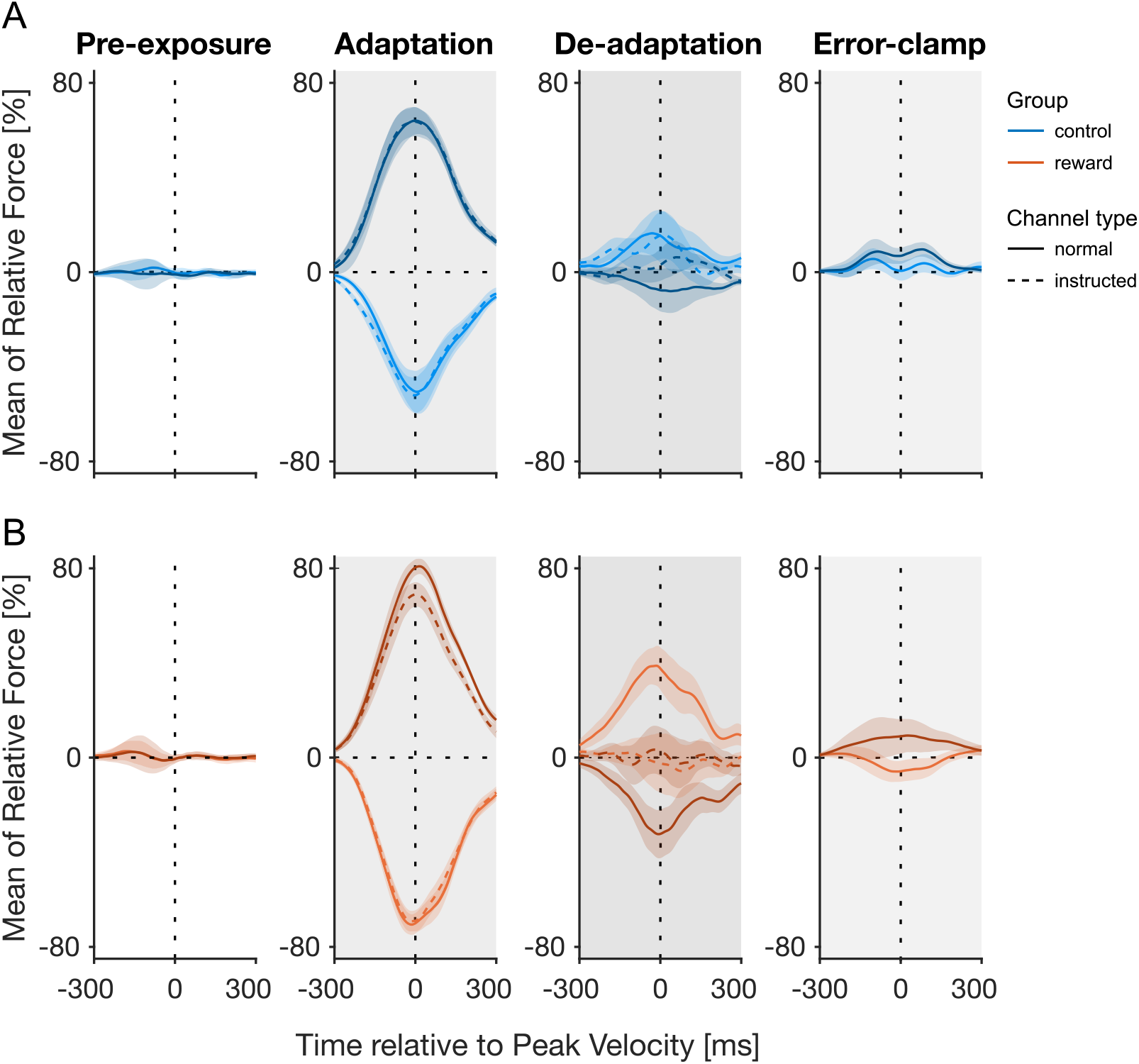
Profiles of predictive force. A. Force profiles on the normal (solid line) and instructed (dashed line) channel trials for the control group as a function of movement time in the pre-exposure phase (all 5 blocks of trials), adaptation (last 5 blocks of trials), de-adaptation (last 3 blocks of trials) and error-clamp (all 5 blocks of trials). The force values are shown as a percentage of perfect force compensation and aligned to peak velocity. The two cues are represented by the dark and light colored lines. B. Force profiles for the reward group.

In the adaptation phase, each contextual cue was associated with one of two curl force fields (clockwise and counterclockwise). Initial trials exhibited large kinematic errors in the lateral direction depending on the force field (cue 1 and cue 2, Fig 4A-B, light and dark colors), with a difference between groups (post-hoc comparison: p=0.019). The early stage of adaptation, being composed of the first five blocks of trials (similar length as the pre-exposure phase, 170 trials) allows us to assess the early effects of the reward. The difference between the control and reward group reflects an effect of the reward already from the early stage of the adaptation. Over the adaptation phase, the kinematic error decreased to 1.8 *±*0.4 cm (absolute average across cues) for the control group (Fig 4A, blue) and to 0.8*±*0.1 cm for the reward group (Fig 4B, orange), with no difference between the two groups (Fig 4C, post-hoc comparison: p=0.253). Additionally, the relative force compensation (solid blue and orange lines, Fig 4D-E) increased up to 56.3*±*0.1% (control) and 75.0*±*0.0% (reward), but with no difference in the levels of adaptation between the groups (Fig 4F, p=0.284). Importantly, participants adapted to both opposing cues simultaneously, which can be seen by the large force compensation in opposite directions at the end of the adaptation phase (Fig 4D-E, light and dark colors), and the velocity-like bell-shape curves of the relative force profiles (Fig 5, adaptation phase). These results indicate that the cues (opposing workspace visual location) allowed dual-adaptation to occur with appropriate adaptation to the two opposing velocity-dependent force fields.

In the following de-adaptation phase, participants experienced initial high lateral errors in the opposite directions for each cue association. However, they were able to reduce their error quickly as their total force compensation decreased back towards pre-exposure level. Overall, the reward group had a much smaller level of kinematic error at the end of the de-adaptation phase (post-hoc comparison: p<0.001). By the end of the de-adaptation phase, both groups had started adapting to the opposite force field with final de-adaptation levels of force compensation of around-15.3*±*4.4% for control participants and-48.6*±*10.6% for reward participants. These results are again supported by the force profiles (Fig 5) showing that the mean relative force for both cues flipped signs at the end of the de-adaptation phase, with a higher absolute peak force value for the reward group. Our experimental design required participants to de-adapt back to the pre-exposure level minimum before switching into the error-clamp phase. Therefore, we expected no differences in force compensation between the groups at the end of this stage, but we actually found one (p<0.001). All control participants did not trigger this switch and therefore performed the maximum amount of blocks of trials in the de-adaptation phase (340 trials). This does not necessarily mean that they reached the expected de-adaptation level and therefore could still be higher than needed. In contrast, many reward participants (9/14) reached this requirement and therefore switched earlier into the error-clamp phase (mean of 235.6*±*29.7 trials), reaching the expected de-adaptation level. Therefore, the difference in force compensation between control and reward participants would be justified. Additionally, we could find a difference in average de-adaptation time between groups, where reward participants de-adapted faster than control participants (t_26_=3.5154, p=0.002).

In the final error-clamp phase, channel trials clamped the lateral error to zero in order to assess the remaining adaptation through the presence of spontaneous recovery [45, 57]. No difference in force compensation between control and reward participants was found (post-hoc comparison: p=1.000). Although the force profiles (Fig 5) show traces of the initially learned force field for the reward group, there was no evidence for spontaneous recovery in either group. Specifically we calculated the difference in force compensation between both cues, and found no deviation from zero for both the control (one-sample *t* test: t_13_=1.583, p=0.137) and reward (one-sample Wilcoxon test, V=80.000, p=0.091) groups.

Overall, no statistical differences in force compensation are seen between he two groups. However, we observe a small qualitative difference in force compensation (Fig 4C) and force profiles (Fig 5) between the two groups at the end of adaptation, that may continue in the following phases. To understand better the discrepancy between the statistical results and the qualitative observations, we examined the individual data in force compensation (Fig 6). In the individual adaptation results, we see a larger variability across cues and participants in the control group compared to the reward group. Instead, the reward participants display more consistent end-level of adaptation. In this case, it is possible that the results tend towards a difference in end-level adaptation, but that the high variability across participants masks any statistical support.

**Figure 6.**
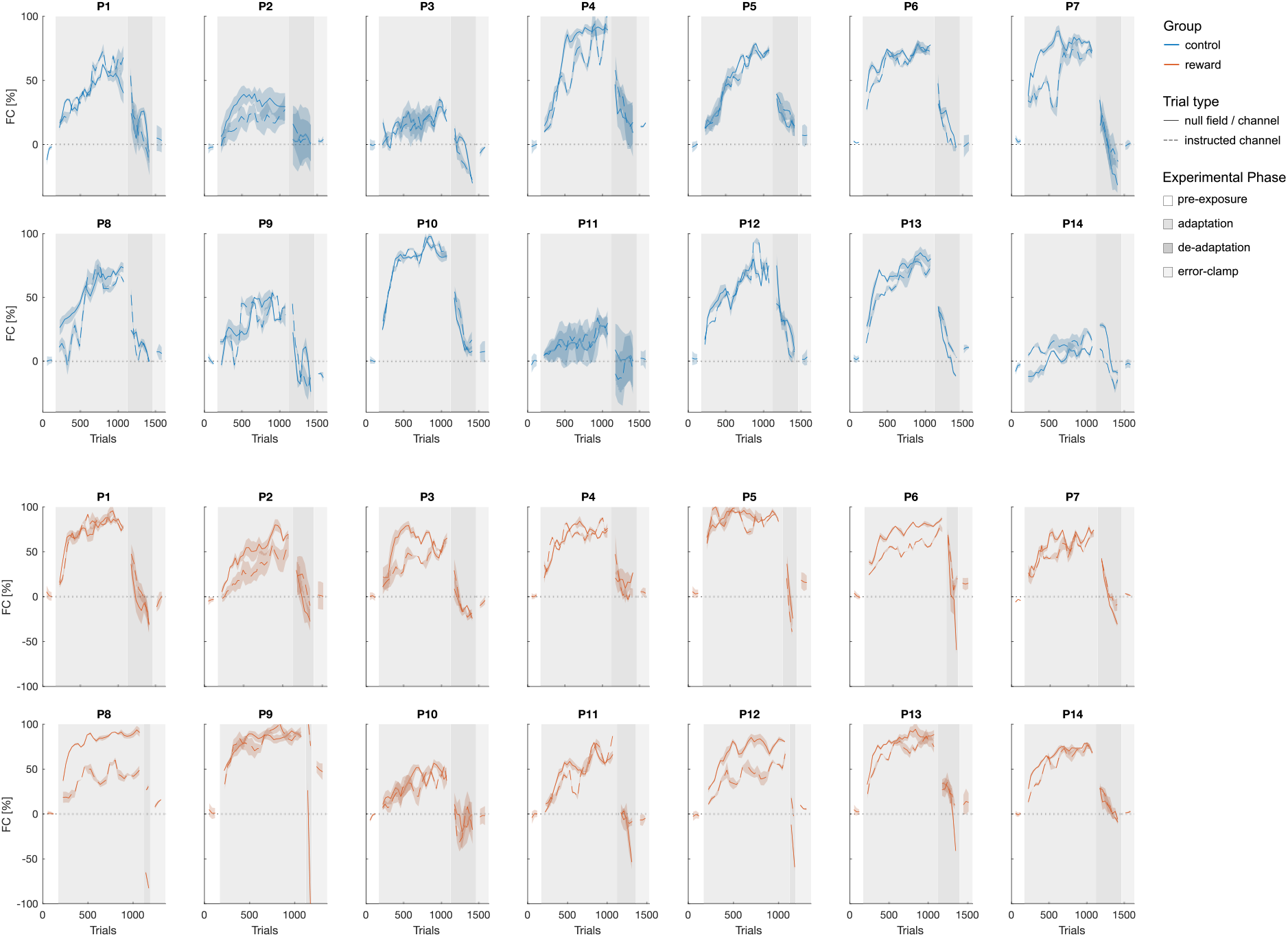
Temporal pattern of adaptation for individual participants. Mean force compensation (solid line) and standard error of the mean (shaded region) across cues for both the control (blue) and reward (orange) groups, throughout the pre-exposure (white), adaptation (grey), de-adaptation (dark grey) and error-clamp (light grey) phases. The mean block values are smoothed using a moving average with a window of 3 blocks.

### Implicit and Explicit Contributions

To further study the effect of reward on adaptation, we dissociated the implicit contribution compared to the total adaptation through the use of instructed channel trials [37, 38], where participants were instructed that the forces had been removed. The concept is that the force compensation observed in instructed trials represents the implicit component of adaptation. Consequently, we assume that the difference between this implicit level of adaptation and the overall level of adaptation, measured on channel trials, reveals the explicit strategies used by participants to improve their performance.

In the control group (Fig 4, 5 blue color), we observed no difference at the end of the adaptation phase in force compensation between channel and instructed channel trials (V=54.000, p=0.952). Indeed these measures were similar throughout the adaptation phase suggesting that most of the adaptation in the control group was implicit in nature. In contrast, in the reward group (Fig 4E-F), we found evidence for a larger force compensation on channel trials than on instructed channel trials (t_13_=2.296, p=0.039). We therefore find evidence for the use of explicit strategies at the end of adaptation in the reward group, while none for the control group. One notable effect is that the rate and amount of implicit adaptation is almost identical between the reward and control groups across the adaptation phase (Fig 4F, 5), suggesting that reward had little, or no, influence on the implicit processes. The de-adaptation phase shows similar results, with no difference between total and implicit adaptation for the control group (V=33.000, p=0.241), but a difference for the reward group (V=20.000, p=0.042). Additionally, we find no statistical difference when we compare the implicit adaptation of the reward group with the implicit (V=52.000, p=1.000) and the total adaptation (V=48.000, p=0.808) of the control group. These results suggest that most if not all of the differences in adaptation between the control and the reward group comes from explicit strategies.

Interestingly in the reward group, while only a small difference is found at the end of adaptation in the total adaptation (normal channel trials) compared to the implicit adaptation (instructed channel trials), reflecting the presence of explicit strategies,a larger difference is observed in the earlier stages of learning. In order to examine whether this might reflect a stronger explicit component, and investigate the effect of reward on the implicit adaptation, we performed further analysis on both the adaptation rate (exponential fit) and the learning and retention rates (multi-rate model).

### Adaptation Rate

We fitted an exponential function to the force compensation data during the adaptation phase to assess the rate of adaptation during the initial exposure. The parameters were estimated using a leave-two out cross-validation sampling (see Methods), providing a total of 91 estimates for each parameter. This resulted in the parameter distributions for both the trial adaptation rate and asymptote for the control (C) and reward (R) groups, and for the normal (C and R)) and instructed (Ci and Ri) channel trials (Fig. 7). The best fit exponential function (7A) and estimated parameters for the asymptote of adaptation (7B) and the trial adaption rate (7C) show clear differences between the control and reward group. In particular, similar to the final adaptation level for the force compensation data, the asymptote is larger for the total reward condition than the other three conditions, although there are differences between all conditions (Fig. 7B). However, the exponential fits allows us to examine the rate of adaptation across all four conditions. We find a faster adaptation rate for the reward compared to the control condition (Fig. 7C) for both normal (R: 0.008,C: 0.0053) and instructed channel trials (Ri: 0.0057,Ci: 0.0033). Furthermore, differences in the adaptation rate between normal and instructed channel trials are observable in the reward (R: 0.008 against Ri: 0.0057) and control groups (C: 0.0053 against Ci: 0.0033). Overall, these results suggest that reward increases the trial by trial adaptation rate compared to the control condition. Additionally, the differences for both groups between the normal and instructed channel trials suggest that an explicit component is present in the overall adaptation to novel force field, that might be increased through the presence of reward. To assess these results in more detail, we fitted the entire experimental data with a multi-rate model for deeper analysis.

**Figure 7.**
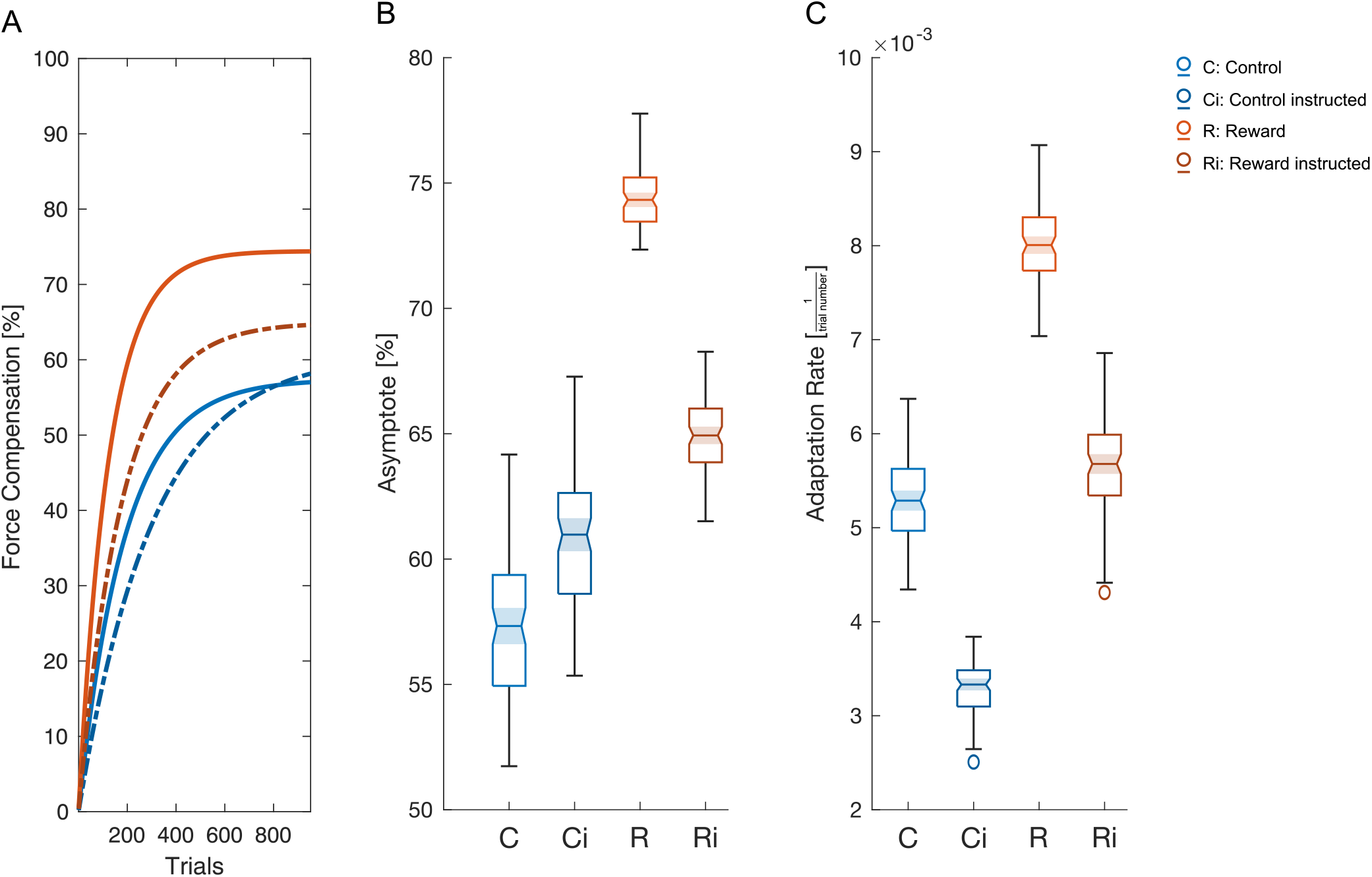
Rate of adaptation to the novel dynamics in the adaptation phase. A. Best fit adaptation curve using the mean parameters across fits to the exponential function for the control (blue) and reward (orange) groups. Solid lines indicate the fits for the channel trials and dotted lines indicate the fits for the instructed channel trials. B. The best-fit parameters of the asymptote of adaptation. Parameters are estimated separately for the instructed and normal channel trials. Model parameters were obtained using leave-two out cross-validation sampling method, which provided 91 estimates of each parameter. Parameters estimates are plotted using boxchart in Matlab, where the line indicates the mean, the shaded notch indicates the 95% confidence intervals, the upper and lower edges of the box contain the upper and lower quartiles, the whiskers contain the non-outlier maximum and minimum, and any outliers are indicated with small circles. If the shaded notch regions do not overlap, then the parameters have different medians at the 5% significance level. C. The best-fit parameters of the adaptation rate (trial constant of adaptation).

### Differences in multi-rate model components

To examine the underlying processes of general adaptation of novel force field and more specifically to dual-adaptation, we used a triple-rate model with a weighted contextual cue switch parameter (Switch) [45]. We therefore assessed the difference in retention (parameter A) and learning (parameter B) rates between the two groups and how they behave over the experiment, as well as how contexts are dissociated through contextual cues (Switch parameter). This model was fitted independently to the force compensation on instructed and non-instructed channel trials in order to separate the implicit and explicit components of adaptation. All parameters were estimated using a leave-two out cross-validation sampling (see Methods), providing 91 parameter estimates of the fast (f), slow (s) and ultra-slow (us) processes for each of the four groups C (Control), Ci (Control instructed), R (Reward), Ri (Reward instructed). These parameter estimates and their medians, displayed in figure 8, show differences across both the groups and channel trial types.

**Figure 8.**
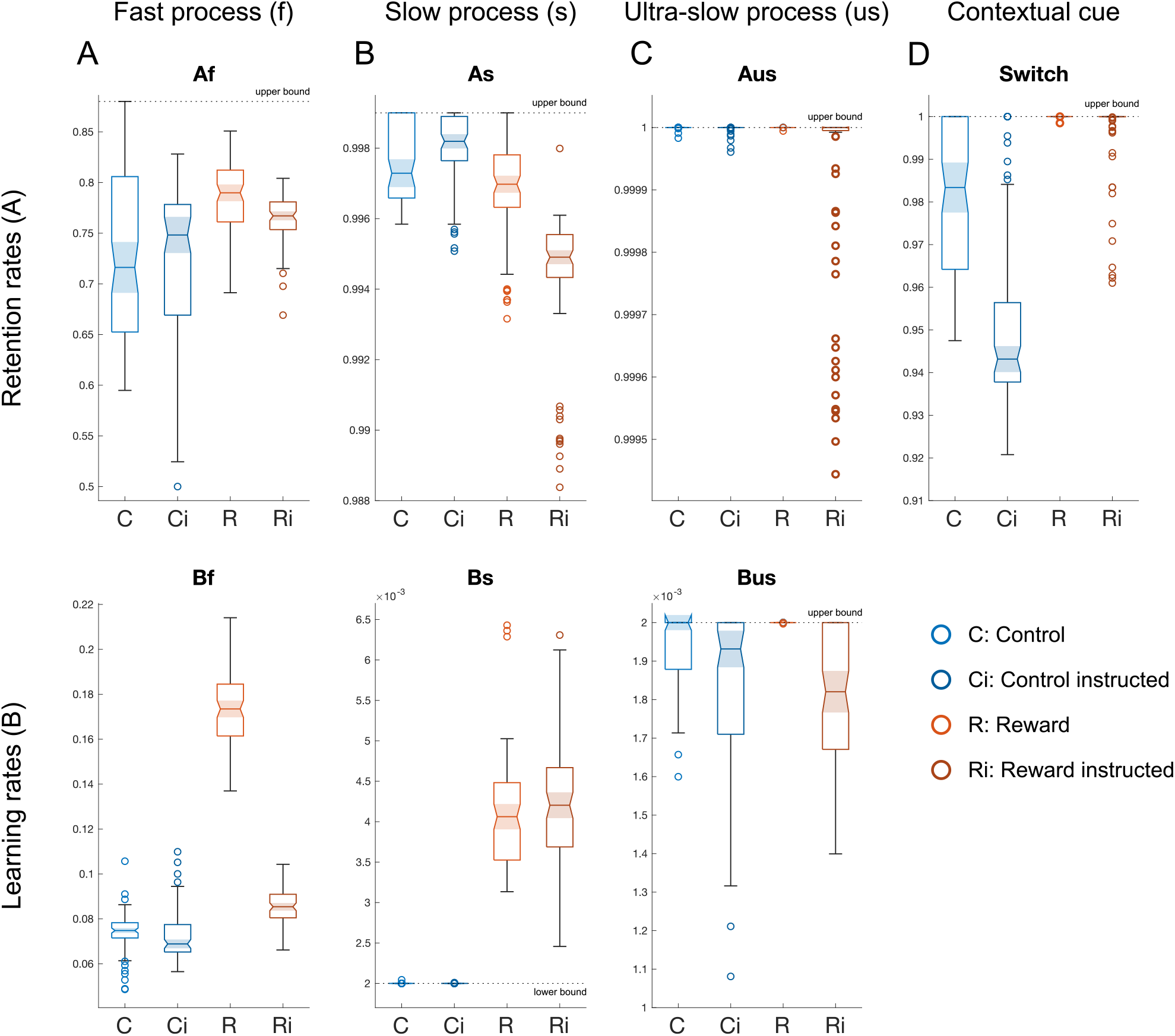
Best-fit triple-rate model parameters. Retention rate (upper row) and learning rate (lower row) parameters fitted to the weighted triple-rate model, containing a fast (A), slow (B), super slow (C) and weighted-switch (D) parameter. The upper or lower bounds (black dotted lines) are displayed when they are contained within the figure axis limits. Parameters are estimated separately for the instructed and normal channel trials. Model parameters were obtained using leave-two out cross-validation sampling, which provided 91 estimates of each parameter. Parameters estimates are plotted using boxchart in Matlab, where the line indicates the mean, the shaded notch indicates the 95% confidence intervals, the upper and lower edges of the box contain the upper and lower quartiles, the whiskers contain the non-outlier maximum and minimum, and any outliers are indicated with small circles. If the shaded notch regions do not overlap, then the parameters have different medians at the 5% significance level.

The fast process (Figure 8A) shows clear differences between the control and reward groups in normal and instructed channel trials for both retention (H(3)=51.1, p<0.0001) and learning rates (H(3)=255.7, p<0.0001). There was a significant difference in the retention rates between reward and control groups for both the non-instructed (Af, C and R, p<0.0001) and instructed channel (Af, Ci and Ri, p=0.0271), as well as within the reward group (R and Ri, p=0.0003) but not within the control group (Af, C and Ci, p=0.3112). For the learning rates (Bf), we found a difference in normal trials between the reward and control group (Bf, C and R, p<0.0001) and the reward instructed and control instructed group (Bf, Ci and Ri, p<0.0001). There was also evidence for higher learning rates in normal compared to instructed trials in the reward (Bf, R and Ri, p<0.0001) but not the control group (Bf, C and Ci, p=0.5195). Importantly, the difference of the learning rates between the instructed and non-instructed trials is very large for the reward (0.0882), which strongly suggests a great use of explicit strategies in the reward group, which might also account for the higher early adaptation (Figure 3). These results suggest that the use of reward increased adaptation through both learning and retention rates of the fast process. Both the retention and learning rates are improved implicitly and explicitly. However, the difference in learning rate’s improvement between reward and control is significantly greater than for the implicit. This suggests that the learning rate is mainly enhanced through explicit strategies, with a small increase in the implicit component, while the retention rate seems to increase lightly implicitly only, with the presence of an explicit component independently from the use of reward.

For the slow process (Figure 8B), we also found differences across groups for both the retention (As, H(3)=189.37, p<0.0001) and learning rates (Bs, H(3)=273.69, p<0.0001). In the retention rate, there is a difference between the reward and control for both normal (As, C and R, p=0.0003) and instructed trials (As, Ci and Ri, p<0.0001). The control group shows no difference between trial types (As, C and Ci, p=0.9995). However, the reward group displays a strong difference between the normal and instructed trial (As, R and Ri, p<0.0001), suggesting an involvement of explicit strategies. For the learning rates (Bs), we find a clear difference only between reward and control conditions for both the normal (Bs, p<0.0001) and instructed (Bs, p<0.0001) trials. However, there are no differences in the learning rates between the normal and instructed trials for either the control (Bs, C and Ci, p=0.7900) or reward groups (Bs, R and Ri, p=0.8717). This suggests that reward only influences the slow learning rate through an increase in implicit adaptation.

Parameters of the ultra-slow process showed evidence of different learning (Bus, H(3)=94.4, p<0.0001) and retention (Aus, H(3)=37.13, p<0.0001) rates across groups (Fig 8C). Although the retention rate had a main effect, we did not find any difference between the reward and control groups for both normal and instructed trials (Aus, C and R, p= 1.0000, Ci and Ri, p=0.3727). No effect was found between trial types for both control and reward groups (Aus, C and Ci, R and Ri, decimal-limit reached). Similarly, although we found a significant main effect of the Kruskal-Wallis test for the learning rate, we found no significant difference in learning rates between groups for instructed trials (Bus, Ci and Ri, p=0.1409) and normal trials (Bus, C and R, decimal-limit reached). While no differences were present between trial types for the control group (Bus, C and Ci, p=0.0530), we found a difference between normal and instructed trials in the reward group (Bus, R and Ri, p<0.0001). While the differences are small, the lower learning rates of instructed trials parameters in both control and reward groups suggest a possible role of explicit strategies.

Contextual cue switch parameter estimates (Fig 8D) also show differences across groups and trial types (Switch, H(3)=188.97, p<0.0001). First, the reward group shows an increase from the control group regardless of trial type (Switch, C and R, p<0.0001; Ci and Ri, p<0.0001). Second, in the control group, the normal trials display higher values than their respective instructed trials (Switch, C and Ci, p<0.0001) but not in the reward group (Switch, R and Ri, decimal-limit reached). These results suggest two main findings. First, the presence of reward increases the weight of the contextual cue switch. Second, it appears that a small, but existent, explicit strategy contributes to the switching between two different cues. However, it is important to note that these small differences include values close to 1. This may suggest little overall impact on motor adaptation when we consider that the range of the switch parameter is from 0.5 (which would be a 50% estimation of the adequate motor memory to update for both opposing contexts) to 1 (100% certainty for the motor memory to update). To evaluate the impact of the differences in the switch parameters, we simulated how adaptation would be impacted by different rates of this parameter in our experimental design (Fig 9). These results clearly show differences between the simulated rates, and therefore between the different groups associated with these rates (R*_c_* and R*i_c_*= 1, C*_c_* = 0.98, C*i_c_*= 0.95). While the differences between the switch parameter are little, they can produce large effects on the overall adaptation.

**Figure 9.**
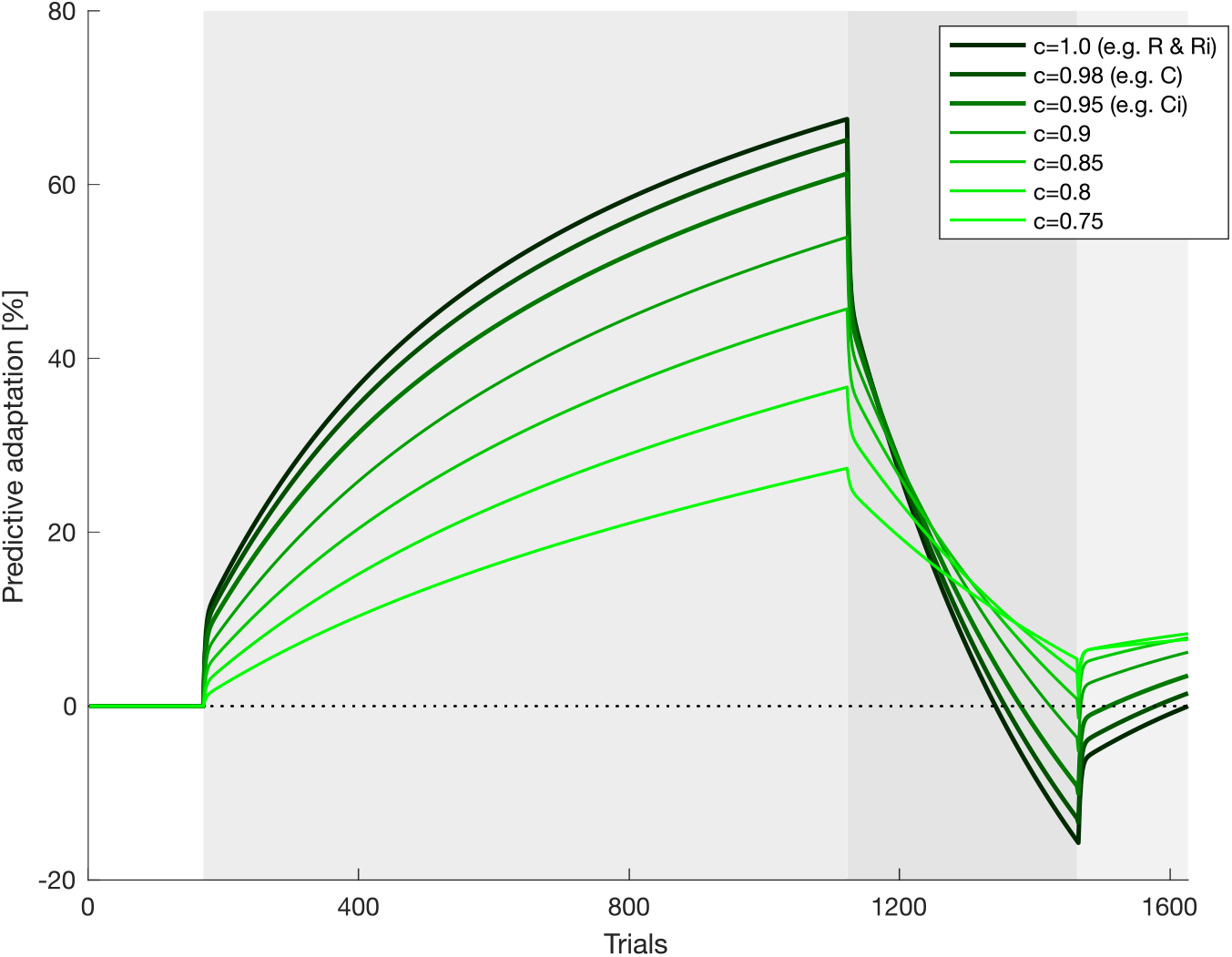
Effect of different contextual cue switch parameters. Simulation of the force compensation over an adaptation, de-adaptation, error-clamp experiment with two different cues. The weighted triple-rate model was simulated with seven different contextual cue switch parameter values (0.75-1). All other parameters were kept constant, using the median of fitted learning and retention parameters for the control condition. The background areas represent the pre-exposure (white), adaptation (grey), de-adaptation (dark grey) and error-clamp (light grey) phases. It is clear that even small differences in the weighting parameter, such as found in the control group, can produce large differences in the overall adaptation.

In our experimental design, we chose a contextual cue known to be very effective in distinguishing two opposing force fields [43, 45]. According to a previous study [38], the use of this effective cue appears to primarily drive implicit learning. Therefore it is possible that any explicit strategy effect on the weight of the contextual cue switch might be hard to distinguish with this type of cue. It would be necessary to further investigate this switch parameter using other types of cues.

In order to ensure that these findings are not specific only to the triple-rate model, we performed an identical experimental fit to the weighted switch dual-rate model (Fig 10). The general pattern of variation in parameters across the four groups of trials exhibits some similarities. Although differences are found in all parameters, one consistent finding is that again reward contributes to an increase in both the fast and slow learning rates (except for the slow process in the retention rate), with evidence that an explicit component is responsible for these differences in the learning rates.

**Figure 10.**
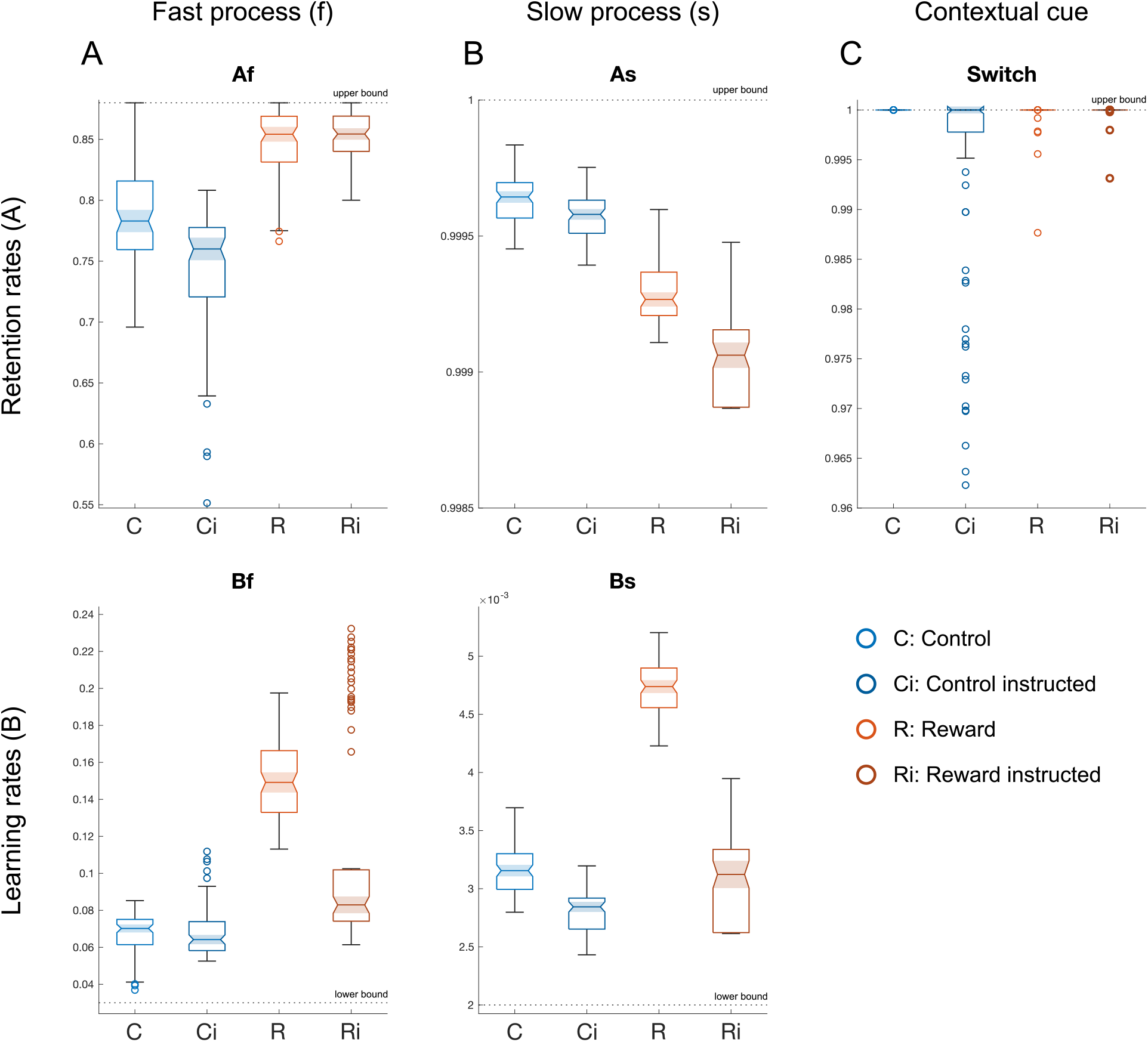
Best-fit dual-rate model parameters. Retention rate (upper row) and learning rate (lower row) parameters fitted to the weighted dual-rate model, containing a fast (A), slow (B) and weighted-switch (C) parameter. The upper or lower bounds (black dotted lines) are displayed when they are contained within the figure axis limits. Parameters are estimated separately for the instructed and normal channel trials. Model parameters were obtained using leave-two out cross-validation sampling, which provided 91 estimates of each parameter. Parameters estimates are plotted using boxchart in Matlab, where the line indicates the mean, the shaded notch indicates the 95% confidence intervals, the upper and lower edges of the box contain the upper and lower quartiles, the whiskers contain the non-outlier maximum and minimum, and any outliers are indicated with small circles. If the shaded notch regions do not overlap, then the parameters have different medians at the 5% significance level.

Over all timescales, while small differences were present in the retention rates, strong effects are seen in the learning rates. Indeed, reward increased the rate of learning of both fast and slow processes. While this improvement arises from implicit mechanisms for the slow process, strong explicit strategies emerged in the fast process. These explicit strategies were also apparent in the contextual cue switch, an important parameter for dual-adaptation, which relies on human’s capacity to dissociate environmental contexts.

Finally, we took the median parameters fit across all participants and conditions and simulated our experiments (Fig 11). These parameters are able to capture all the major effects found in the experimental data (Fig 12). Specifically we find a slightly faster and higher adaptation in the reward normal trials compared to the other three groups (control normal, control instructed and reward instructed trials). They also capture the faster de-adaptation in the reward group and slightly higher rebound of the predictive forces in the error-clamp phase. Overall, this shows that the parameters estimates obtained are able to re-capture most of the key elements of the experimental results (Fig 4D-F).

**Figure 11.**
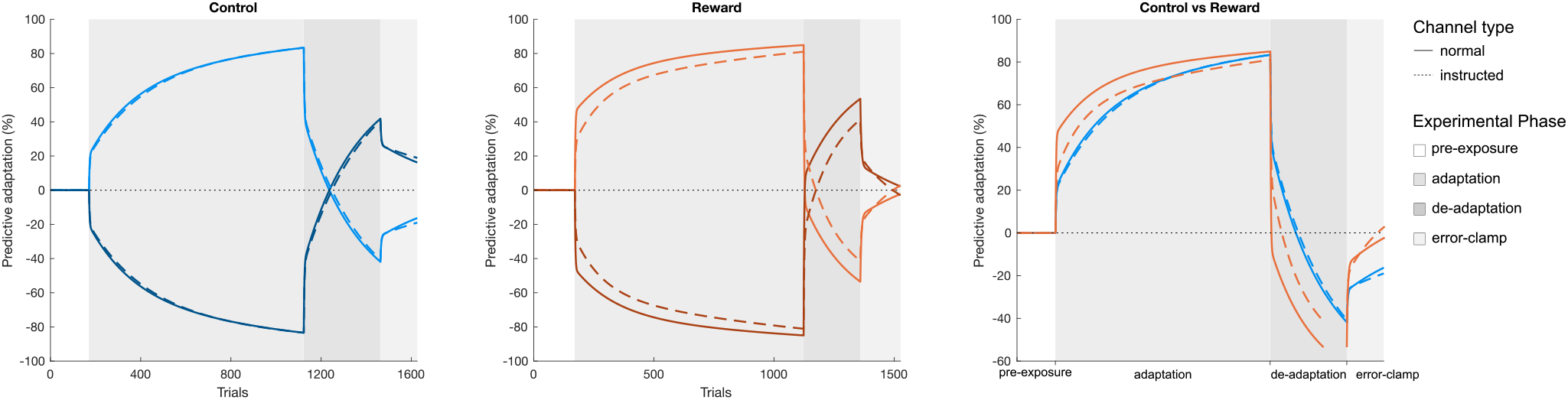
Predictive adaptation of our experimental design using the best-fit parameters. Simulation of the force compensation on normal (solid line) and instructed (dashed line) channel trials for the control (blue) and reward (orange) groups. The experiment was simulated by the weighted triple-rate model using the median of fitted parameters for the learning and retention rates.

**Figure 12.**
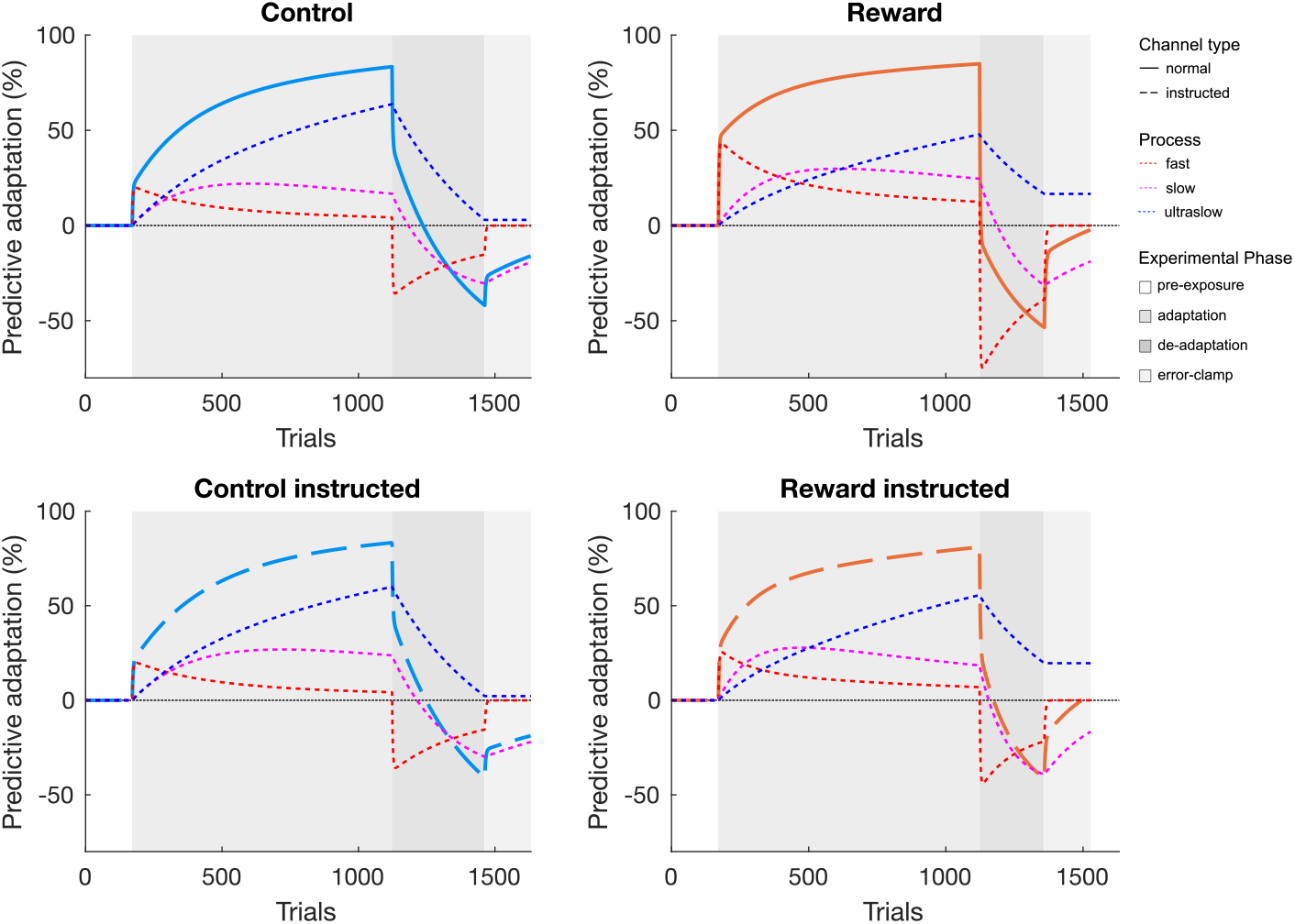
Predictive adaptation of our experimental design using the best-fit parameters displaying its three timescales. Simulation of the force compensation on normal (solid line) and instructed (dashed line) channel trials for the control (blue) and reward (orange) groups displaying the fast, slow and ultra-slow processes (red, magenta and dark blue dotted lines, respectively). The experiment was simulated by the weighted triple-rate model using the median of fitted parameters for the learning and retention rates. As both cues are equivalent in profile except for a reversed sign, only one cue is displayed for clarity.

## Discussion

The goal of this study was to investigate the influence of reward on dual-adaptation, and more specifically on the different components of adaptation. A reward group and a control group simultaneously adapted to two opposing force fields cued by visual feedback location. We found an increase in success level for the reward group suggesting overall straighter reaching movements, with differences between control and reward groups in late adaptation and de-adaptation for both bias and variability. Participants of both groups were able to adapt independently to the two opposing force fields, reaching a roughly similar final adaptation level at the end of adaptation, with some evidence for explicit strategies contributing to slightly higher and faster adaptation in the reward group. While the reward group displayed a faster de-adaptation than the control group, we found no difference in the error-clamp phase between groups. To examine how reward affects different adaptive processes, we fitted a weighted triple-rate adaptation model to the experimental force compensation data on normal and instructed trials. The reward group showed clear differences on the fast and slow timescales of adaptation but no effect on the slowest timescale: the ultra-slow process. Differences in the model parameters on the instructed channels and between instructed and non-instructed channels demonstrated that reward influenced adaptation through both implicit and explicit adaptive processes. Reward drove changes in implicit adaptation, producing small increases in both the fast learning and retention rates, but strong increases in the slow learning rate. In addition reward produced a strong explicit increase in the learning rate of the fast process. Although the contextual cue switch parameter (for our experimental design) is primarily implicit in nature according to the control group results, it appears that this is enhanced by an explicit process, that appears to increase in the presence of reward. Supporting previous findings [38], we found no major contribution of explicit adaptation to the control group adaptation processes, suggesting that learning of novel dynamics is primarily implicit in nature in the absence of rewards or punishments. In our experiment, we designed the reward to be dependent on the participants’ horizontal kinematic error (maximum perpendicular error). Therefore, in order to increase reward, participants needed to execute straighter movements. Indeed, we found straighter trajectory profiles from the reward group compared with the control group. However, not only were the participants straighter and reduced their horizontal kinematic error faster, but they also became more consistent in their movement, with a reduced trial-by-trial variability in these trajectories as seen previously [13]. Previous studies have considered variability as an implicit form of motor exploration during periods of low success in search of a more rewarding state [6, 67]. In this framework, movement variability would decrease in periods of high reward probability, as the need of changing their current successful motor command decreases [68]. This suggests that participants achieve a higher state certainty and therefore perform more consistent movement [4, 69]. Our present results are consistent with this idea that variability in motor commands is partly driven by the history of reward. However, it has also been shown that reward can increase accuracy and decrease variability through increased muscle co-contraction and limb stiffness [70], which has also been shown to increase the speed of adaptation to novel force fields [71]. Indeed, an increased muscle co-contraction is also consistent with our results, which show a greater difference in end-level of kinematic error between reward and control participants compared to the difference in end-level force compensation. While we did not record muscle (EMG) activity, we would predict that there was a higher level of muscle co-contraction in the reward group compared to the control.

Statistically, participants of both groups exhibited similar levels of force compensation and kinematic error at the end of adaptation, unlike a previous study of force field adaptation which showed different final adaptation levels for reward and control conditions [14]. There are possible reasons for these differences. First their study examined older stroke survivors, whereas our study focused on young healthy adults. Second, their design had a shorter adaptation phase of 350 trials, compared with our 952 trials, and only examined adaptation to a single force field whereas we examined simultaneous adaptation to two opposing force fields. It is possible therefore that the final levels would have eventually came to the same level for both control and reward groups if a longer adaptation phase was provided. Third their study examined kinematic error reduction rather than force compensation, which means that different levels of final adaptation might also reflect differences in limb stiffness contributions to error reduction [72, 73]. However, it is important to note that we do find evidence for differences across conditions with the estimates of the asymptote from the exponential fits. Indeed the asymptote estimates are similar to participants’ end-level of force compensation, with evidence for clear differences between each group. Additionally, the qualitative observation of the end-level of force compensation also shows a small difference between groups. With the difference between groups shown by the exponential fit, it also seems plausible that the reward group achieved a slightly higher asymptotic performance by the end of the adaptation phase, similar to the previous work. In this case, it is possible that a high inter-individual variability in our current results hides a real difference of adaptation between groups. Finally, we do find similarities with the study of Quattrocchi and collaborators [14] in the beginning of the adaptation phase, as well as in the de-adaptation phase. These both reflect early stages of adaptation, for example towards the second opposing environment for the de-adaptation phase, due to the short trial number in this phase. In both stages, we found that reward produced faster adaptation, or de-adaptation,in the early parts of the adaptation process, a claim that has been made by Quattrocchi and collaborators [14]. Overall our findings are consistent with their study in that reward improves early adaptation to novel dynamics.

In order to quantify differences in implicit and explicit adaptation between the reward and control groups we assessed the asymptote and rate of adaptation, and further the learning and retention rates of adaptation for each group to objectively compare their differences. The exponential fit showed a clear use of explicit strategies for both control and reward groups, a higher adaptation rate for the reward group, and evidence for stronger use of explicit strategies in the final level of adaptation (asymptote). To examine the specific effects in more detail, we fitted a triple-rate model of adaptation including a fast, slow and ultra-slow process to the data of each group. These results allowed us to compare the previously analysed processes at the end of adaptation, to earlier adaptation stages. Although it depends on the exact parameters, we can imagine that the contributions of each of the different processes of the triple-rate model analysis are more active at different times within the experiment (Fig 12). In this case, the fast process, highly sensitive to movement errors, mainly drives the adaptation in very initial stages of force field exposure and is followed by the slow process which contributes most in the middle of adaptation, when participants’ error has been reduced. Finally, the ultra-slow process takes over in the late exposure. The reward group had higher fast and slow learning rates, which explains the faster reduction in error for the reward group in the early stages of adaptation. Similarly, the higher fast and slow learning rates strongly influence the de-adaptation of reward group, who exhibited significantly shorter numbers of trials to reach a similar level of de-adaptation. This clear difference in learning rates are in line with previous studies that claimed an increase of learning rate in rewarded contexts [12, 13].

This present study aimed to assess how reward influences implicit and explicit contributions to motor adaptation. While reward affects the slow learning process primarily implicitly, it strongly affects fast learning through the use of explicit strategies, with a small contribution of the implicit component. Several studies have suggested the use of explicit strategies as a mechanism to improve motor performance in rewarded states [74, 75]. One theory relies on the fact that explicit strategies mainly drive early adaptation in order to reduce systematic errors, as a compensation for the slower implicit adaptation [76–80] activated through cerebellar brain areas [4, 28, 81–84]. In addition, while the implicit component increases over time, the use of explicit strategies reduces, inducing a total adaptation relying more on the implicit component of adaptation [85, 86]. Our current results show no explicit component in the slow and ultra-slow process learning rate and a high implicit component in the slow process. Additionally, the de-adaptation phase reveals that reward participants rely only on explicit strategies in order to de-adapt faster than the control participants, with a similar level of implicit adaptation for the reward and control group, and for total adaptation for the control group. As the implicit adaptation appears at the same level at the end of the de-adaptation phase as well as in the error-clamp phase, it is realistic to suggest that the retention is mainly driven by the implicit component through slower timescales. Furthermore, this implicit-only ultra-slow process did not show an enhancement in motor performance in the reward group compared to the control group, explaining why we find similar final adaptation levels in both groups. This additionally supports the idea that explicit strategies produce rapid improvements in performance involving prefrontal and pre-motor cortex [23, 24, 26, 28, 30, 81, 87–89], and are highly sensitive to error [90–94]. All together, these results support the theory that explicit components lay more in the fast processes whereas implicit components lay more on the slower timescales. In our experiments the reward signal primarily drives these explicit strategies, and therefore increases the speed of early adaptation, but has little effect on the final level of adaptation.

It is important to note that the control group did not display any evidence for explicit strategies in dual force field adaptation. This is in contrast to several studies which have linked the fast process with explicit strategies. They suggest that error-driven motor learning uses a fast prefrontal explicit component that disappears as the slower cerebellar implicit component is activated [29, 76, 77]. Our results argue against the possibility that the fast process fully represents the explicit component, at least in dual force field adaptation. However, it is also important to take into account our experimental context in which we used a highly effective or direct cue, that has been shown to drive implicit-only adaptation [38]. Therefore, it is conceivable to imagine this fast process to be highly flexible, using some implicit adaptation when possible, such as in the slow process, but enabling the use of explicit strategies depending on the context. Indeed, the presence of reward showed strong effects on explicit learning both in the fast and slow processes, suggesting that neither the fast nor slow processes can be considered fully implicit or explicit. Indeed, we find clear evidence that reward increases the fast retention rate through implicit processes, both in the triple and dual-rate models. This supports the existence of implicit components at any timescales of adaptation, that would be volatile, or modulated by specific characteristics of the learning environment [95], such as trial-to-trial consistency [35, 96] and the certainty of context estimation [97, 98].

Several studies have suggested that reward primarily affects retention rather than learning rates [15, 17–20]. In contrast to these results, our work shows that reward had a very strong effect on the learning rates of both the slow and fast processes. However, the effect of reward on the retention is much less clear in our study. In the fast process, reward increased both implicit and explicit components of the retention rate. However, the reward produced a decrease in the retention rate in the slow process, and no effect on retention in the ultra-slow process. Taken all together, we could think that, with an improvement in retention in the fast process, reward enhances the overall retention rate. This statement would support literature that showed a main improvement in retention rate by reward [17, 21, 35, 99–102]. However, an increase in the fast retention rate, with no effect on the slower timescales will only speed initial adaptation and have little effect on the final level of adaptation, or even retention of learned motor memories over much longer timescales. It is possible that decreases in the slow retention rate with reward might be related to our specific model of adaptation and constraints on the parameters. However, it is important to note that even the dual-rate model (Fig 10), with no limitations via constraints on the upper bound of the retention rates of the slow process, showed a decrease in the slow retention rate with reward. Another key consideration is that our experimental design did not contain long periods of channel trials where retention could be easily parameterized with the exception of the error-clamp phase. However, our previous study [45] showed that a design including additional long periods of channel trials to estimate retention directly, and a longer error-clamp phase to examine spontaneous recovery, did not show differences from the results using a similar experimental design to the one used in this study. Overall, these results might suggest that reward does not fundamentally influence long-term retention, at least in our experimental context.

This present study demonstrated the presence of both explicit strategies and implicit adaptation for the fast and the slow process. This comes in contrast with multiple studies that have suggested that the fast and slow processes mainly reflect explicit and implicit components of adaptation respectively. It is essential to specify that this work has been done using visuomotor rotation designs only, a design that strongly exploit explicit strategies [29, 32, 77, 78, 103] and show extremely fast adaptation processes. In contrast, force field adaptation appears to be mainly implicit in nature [37, 38] and proceeds at a much slower scale, especially in dual-adaptation design as used in this current study. This makes it difficult to directly contrast the findings for multiple reasons. For example, the fast process in visuomotor adaptation usually refers to a strategic shift in the movement direction, an explicit strategy that may not contribute to force field adaptation at all. Additionally, the limited contribution of implicit adaptation to visuomotor rotation studies may make it much more difficult to detect any reward-based effect on the component. However, interpretation of a reward-based effect on implicit adaptation is more straightforward. A recent study by Sugiyama and colleagues (2023) [10] has suggested that reward directly influences the learning rate to maximize rewards-a meta-learning approach-which explains variations in implicit adaptation through rewards or punishments. While the exact neural location of such a process is unclear, reward has been shown to affect both basal ganglia [104, 105], which plays a role in both implicit and [106–108] and explicit components [109–111], and cerebellar processing, that has also shown an involvement of both implicit [112–115] and explicit components [78, 116–118]. As a whole, we believe that such implicit adaptation is combined with a variety of explicit processes, which could include strategies, increases in co-contraction or other mechanism to reduce the perturbing effects of novel tasks.

All studies which model adaptation through learning rates and retention rates are limited in their ability to fully distinguish between effects on the learning and retention rates. This is because learning rates and retention rates interact in order to model the total behavior, which means that the specific values are dependent upon one another. Thus, a prioritized increase of learning rate would automatically decrease its associated retention rate to fit the overall data, in the absence of additional constraints on the model. To counterbalance this limitation, researchers have applied different constraints to these parameters. It is important to carefully think these upper and lower constraints to keep the processes in time ranges specific to their associated timescale. For example, our results find that reward produces a strong increase in the slow learning rate, which is coupled to a small decrease in its associated slow retention rate. Also, the interactions between our coupled parameters push certain parameters to the bounds, for example with the slow learning rates in the control group. These effects are present in all studies that use such models. Therefore, most studies that model such interactions in the learning and retention, either within or across timescales, include phases within the experimental design in which specific components contribute most to the overall motor output or behavior. For example, de-adaptation phases allow separation within different timescales, and error-clamp phases allow for measurements of the retention rate independent of learning rates. Nevertheless, the interactions across the processes are an inherent limitation of all of these studies and limit the ability to fully capture the exact parameters of the model. Such effects can also explain the differences that each studies finds on the effect of reward on different components of the adaptation process. To explore such limitations, we also fit our experimental data with the dual-rate model. The goal was to examine if most of the parameters vary in a similar pattern, and whether we would observe any difference between the parameters of the triple-rate model that are at the bounds. It is important to note that it is likely to see differences in the specific fits provided by different models, as the constraints are different for each model type. Here we see one major difference between the two models, with the slow learning rate in the instructed channel showing a decrease in the Reward condition, which would suggest that the increase in learning rates with reward are primarily explicit. However, we believe that the triple-rate model is the better overall model for the data, as our previous study [45] found this model best explained data in a similar experimental design using model comparison. Altogether, the similarities in the pattern of interactions between parameters between the dual-rate and the triple-rate support our overall results.

One common way to define the total amount of explicit strategies by taking the difference between the total amount of adaptation and the amount of implicit adaptation [62, 63]. However, this idea that adaptation represents the summation of an implicit and an explicit component has been challenged by several studies showing that implicit and explicit do not always sum up to total behavior [119, 120]. However, as we find only small contributions of the explicit component to the overall adaptation process in our study, even different approaches to separating these components will likely produce similar results. We therefore believe our method to be appropriate for a first approach to assess the impact of reward on the implicit and explicit components over multiple timescales.

One very interesting finding is that explicit strategies contribute to the contextual cue switch parameter, as can be seen clearly for the control group. Here we find that there is a small, but significant increase in the switch parameter that is associated with explicit strategies. Contextual cues can be more or less effective depending on their type [43], where ineffective contextual cues require the presence of explicit strategies to account for dual-adaptation, while effective cues drive implicit adaptation directly [38]. Here we used a visual location contextual cue, chosen as it effectively supports dual-adaptation [38, 43], which led to a very high switch parameter. That is, the use of a highly effective cue, improves the distinction between the states associated with each force field, increasing the certainty of a specific context. In such cases, participants easily estimate the environment, switch appropriately between the conditions, and selectively update the respective motor memory. In our study, we found that the switch parameter was strongly cued implicitly, but with a weak but clear explicit component. In our study, where all switch parameters were very close to one, the small explicit component would contribute little to the overall adaptation. However, dual-adaptation in conditions with less effective cues, which will produce little implicit switching, may exhibit strong explicit contributions to the switch parameter. That is, explicit contributions to switching is likely accessible when needed. It is essential to further test alternative cues to determine the differential contribution of this explicit component to motor memory selection.

The increase in contextual switch parameter for the reward group compared to the control group indicates an additional effect of reward on motor learning. The latter seems to increase the switch weight for both implicit and explicit components. With a stronger or more certain selection of motor memory, we can claim that reward influences the probability of selecting the adequate motor memory to update [35, 121]. We expect this effect of reward to be stronger when paired with less effective contextual cues. Although the current design does not allow us to determine whether reward effects the switch parameter through explicit or implicit mechanisms, we predict that reward would produce a strong explicit contribution. However this would need to be tested under different conditions, with cues that do not produce such strong contextual effects. Overall, we propose that an explicit component might be used in dissociating contexts or interpreting the contextual cues, and would be enhanced through reward.

Overall, we investigated the contribution of reward to implicit and explicit mechanisms of dual-adaptation. Although reward produced similar levels of final adaptation to the control group, we found evidence of faster adaptation to changing dynamics. While control participants primarily adapted from error-based adaptation through sensory prediction errors [23, 27], reward participants benefited from additional reward-based prediction error to execute the task. Error-driven adaptation is an efficient and sufficient process to adequately adapt to simultaneously daily life tasks [5]. However our study highlights how reward modulates this adaptation on different timescales, with faster adaptation towards opposing force fields. Moreover, we could distinguish explicit from implicit components, demonstrating that reward increases explicit strategies to drive this faster learning. This explicit component is likely activated in reward-based adaptation by different neural components, acting independently from error-driven learning [6].

## DATA AVAILABILITY

Source data for this study are now available under the following private link: https://figshare.com/s/85e1f492821a9fe7490c.

